# Shared Rhythm to Shared Vision: Synchronous Marching increases Conformity on Perceptual Decision making

**DOI:** 10.1101/2024.11.21.622221

**Authors:** Manisha Biswas, Marcel Brass

## Abstract

Previous research on synchronous movement rituals have found that it increased prosocial propensity toward the synchronised group and reliance on the group opinions. However, whether basic cognitive processes such as perceptual decision-making are affected by synchronous movement remains unexplored. In this novel virtual reality experiment, we examined whether marching synchronously with a group can induce greater informational conformity on an unrelated perceptual task (forced choice random dot motion). We found higher degree of conformity following synchronised marching during low stimulus information. This finding indicates that synchronised movement induces minimal group membership expressed via increased self-other blurring and conformity. Thus, participation in movement-based rituals has the capacity to change our perception of the world to align more closely with the synchronised group.

## Introduction

Group rituals that incorporate synchronous movement can create a sense of euphoric collective effervescence, a feeling of being “*swept away.... participants experience a force external to them, which seems to be moving them and by which their very nature is transformed*” (1). These synchrony-induced behavioural and cognitive transformations has been the subject of extensive research which consistently demonstrates that it elicits heightened feelings of connectedness, affiliation, and pro-sociality toward synchronised partners (2–6). Interpersonal synchrony serves as a cue for in-group membership from an early age (7,8) and functions as similarity cue (9). By increasing self-other blurring (10–13), it fosters a sense of connectedness and enhances group cohesiveness (14,15).

Cohesion in groups is often expressed via conformity to jointly held group perceptions (16,17). For example, a study by Ross & Levy (2023) showed participants two pictures of presidential inaugurations— from 2009 and 2017—and asked them to identify which had the larger audience. Surprisingly, a large number of respondents selected the incorrect image (with an error rate of 15% among Trump voters compared to 3% among Democrats and non-voters), this illustrates that conforming to the opinion of the political group can sometimes become a stronger motivator than providing the correct answer in decision-making tasks concerned with an objective (and apparent) physical differences. In this case, conformity may have occurred through two possible mechanisms: (1) a deliberative and instrumental process to align with a group’s opinion, or (2) an altering of perception via unconscious and private adoption of group viewpoints (19,20). It has been established by prior research that participation in synchronous activity induces the former kind of deliberative conformity (Dong et al., 2015; Mogan et al., 2017; Wiltermuth, 2012; Wiltermuth, 2012) but the latter kind of conformity which is concerned with changes in basic perceptual decision-making is yet to be investigated in the context of interpersonal synchrony.

Social influence research is concerned with how our social environment impacts our perceptual decision-making, known as informational conformity (24–26). Some researchers argue that social influence permeates and modifies early stages of perception (27–30), by altering early attentional resources and stimulus processing (Germar 2016). Interestingly, conformist tendencies persist even when group members are presented virtually (25,31). For example, Kraemer (2013) successfully replicated Asch’s (1951) perceptual conformity experiment in Second Life, a desktop-computer based virtual world. Furthermore, conformity is amplified in settings where individuals identify with an in-group (33). Using the autokinetic illusion paradigm (Sherif, 1935), in which participants collectively reach estimates regarding an optical illusion (the motion of a light point in a dark room), Abrams et al., (1990) demonstrated that social influence was more pronounced when participants possess a shared identity.

Although interpersonal synchrony has been studied alongside conformity paradigms in the past (4), these investigations have not explored its effects on perceptual decision-making. Dong et al. (2015) conducted a series of experiments showing that participants in the synchronous movement group were more likely to rely on general majority opinion regarding product preferences, this kind of preferential decision making comprises higher cognitive level deliberative processes distinguishing it from perceptual decision making (35). Additionally, another set of studies focus on the relationship between interpersonal synchrony and higher levels of compliance to authority.

Participants in the synchrony condition were more likely to comply with their confederate’s request to administer a noise blast and in another study were more likely to comply with a directive to grind more bugs (Wiltermuth, 2012a; Wiltermuth, 2012b). While these studies demonstrate the far-reaching consequences of participating in synchronous movement, they do not address the fundamental problem of biassed perceptual decision making. If the groups we are aligned with can mould our perception, it is crucial to comprehend which group induction mechanisms can activate these perceptual changes and to carve out the extent of their social influence. In the experiment described below, we focus on shifts in perceptual decision-making as a consequence of synchronised movement induced group identity by directly comparing performance on a random dot motion task alongside synchronised and unsynchronised partners.

In the present immersive virtual reality experiment, participants were exposed to two groups. One marched synchronously, while the other marched asynchronously with the participant and each other. Then participants performed a random dot motion decision-making task alongside each of the two groups. The use of immersive virtual reality (VR) in experimental psychology research, although relatively new, holds promise. It has proven fruitful in research on both synchrony (36,37) and conformity (38–40). We sought to recreate the sensation of synchronous marching with virtual agents as it afforded control and consistency of the synchrony experience across participants (41). By utilising virtual reality (VR), we were able to present life-sized interactants animated with biologically realistic motion, thereby creating an immersive experience of group interaction. We used synchronous marching as our experimental manipulation as it is a widespread ritualistic practice associated with the sensation of collective effervescence (McNeill, 1997; Wiltermuth & Heath, 2009). Furthermore, a recent field study using marching in groups found that it increased cooperation (44). For our perceptual decision making task, we employed the classical random dot motion task (30,45). Previous research has found that it is susceptible to social influence (46). Notably however, the combined exploration of interpersonal synchrony and conformity within a VR experimental paradigm is novel.

## 1. Materials & Methods

This study was pre-registered at https://aspredicted.org/FTL_7WC. The experiment was designed with UXF (47) and run on Unity (http://unity3d.com/unity) using HTC VIVE PRO (https://www.vive.com/de/product/vive-pro/) headset with a frame-rate of 90 fps. Post-task questionnaires were presented on a desktop-computer using Psychopy (48). As pre-registered, a total of 60 participants (Mean age = 26.7; 41 women) were recruited. After the exclusion criteria were applied, 58 participants remained (Mean age = 27.3; 37 women). One participant was removed due to an error in recording of motion data, specifically, the right and left arm were switched by the system at random. Another participant was removed because they lost too many trials through the pre-registered trial exclusion criteria and had several cells of the design missing. The experiment was approved by the ethical review board at the Institute of Psychology, Humboldt University of Berlin and was conducted in accordance with the directives laid out in the Helsinki Declaration of Human Rights.

### 1.2 Design

The design consisted of 3 within subject factors: Synchrony (Marching Synchronously or Asynchronously), Task Difficulty (Easy, Hard, or Ambiguous) and Group Correctness (Correct or Wrong). The synchrony manipulation was blocked and counter-balanced. Participants were placed in a unity scene such that they had a full view of all characters in the scene (see Fig. 1) as well as the white screen used to present the Random Dot Motion task. Each main block consisted of 150 conformity trials, for a total of 300 trials per participant excluding practice trials.

**Figure 1.**
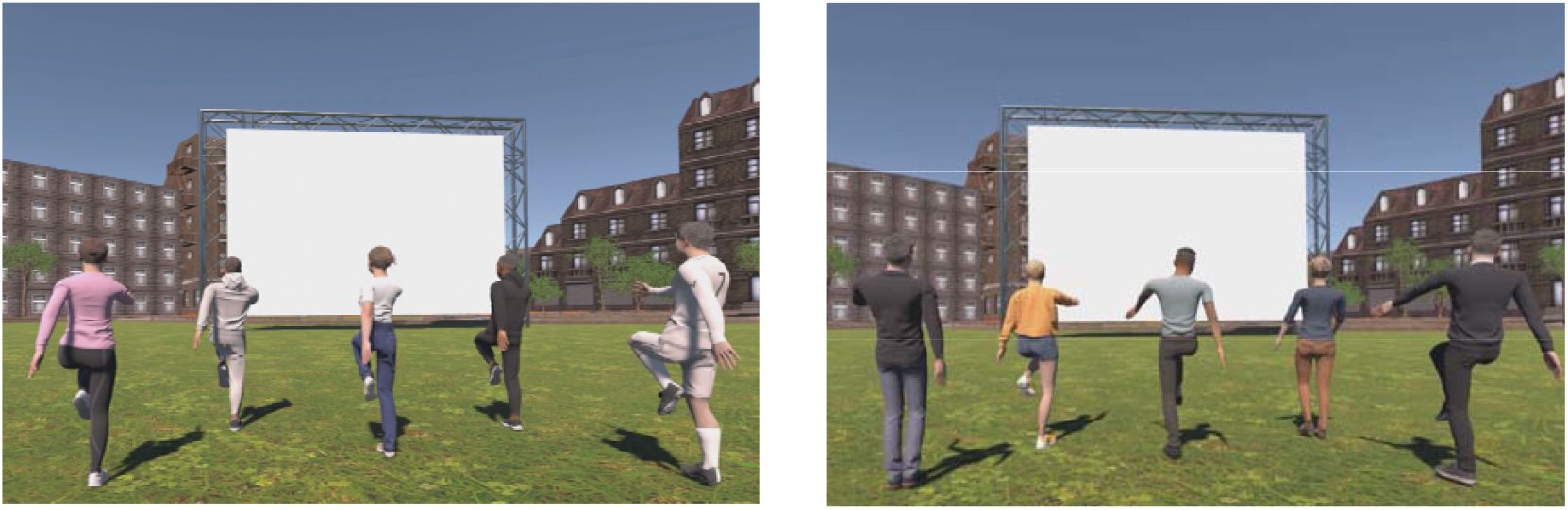
This image depicts two marching manipulations. Left Panel: Group performing synchronous marching Right panel: group performing asynchronous marching.

### 1.3 Marching Manipulation

We manipulated marching behaviour in two virtual groups, presenting participants with one group that marched synchronously and another that marched asynchronously (see Fig. 1). The virtual group that engaged in synchronous or asynchronous marching was counterbalanced. To ensure biologically plausible movements, we utilised 3D full body motion capture produced through the Opti-track system to animate the agents’ marching. Synchronous marching was characterised by a phase-locked and identical movement pattern and participants’ movements were tightly synchronised with the group. In order to achieve this, the group’s marching was timed to a 60-bpm rhythm; participants heard an audio cue timed to the same rhythm and were asked to use this as a reference beat to synchronise their own marching behaviour. Conversely, in the asynchronous condition, movements were decoupled. Participants maintained a 60-bpm marching rhythm with the reference beat while each agent in the group was assigned a different rhythm (ranging from 40 to 80 bpm) accompanied by different movement artefacts derived from various individuals (see left image, Fig. 1). Additionally, the animation assigned to each agent was randomised for each marching block.

### 1.4 Random Dot Motion Conformity task

Following each marching phase, we interleaved perceptual decision making trials. A series of random dot motion trials were presented on a white screen in front of the participant and the virtual group. Approximately 100 dots appeared on the screen, and they moved either to the left or right with varying levels of coherence, depending on the trial difficulty. In easy trials, 30-40% of the dots moved coherently in one direction and in hard trials, 0.1-10% of the dots moved coherently in one direction. For ambiguous trials, the dots exhibited no coherent pattern of movement, making it impossible to provide a correct response. The dots disappeared after 1000 ms, and participants were instructed to raise their hands to indicate the perceived direction of dot movement. The group’s response occurred precisely at 100 ms, ensuring that participants observed the group’s decision before providing their own responses. All members of the group consistently made the same decision and raised their arms simultaneously, as illustrated in Figure 2. Both groups were programmed to display 50% accuracy across the easy and hard conditions.

**Figure 2.**
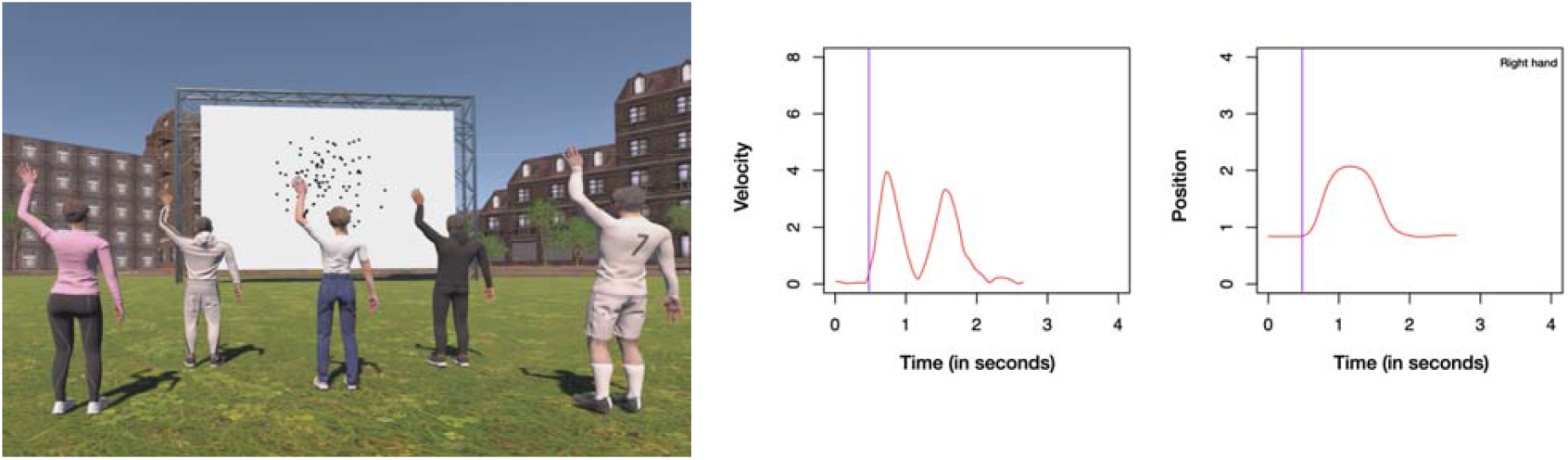
Left Panel: Random Dot Motion/Conformity Task. Right Panel: An example trial showing a raised hand for manual inspection, with velocity and position plotted over time. The purple line indicates the calculation of Reaction Time.

### 1.5 Post-Task Questionnaires

After each block, participants completed a questionnaire to evaluate their subjective experience. They viewed a picture of the group they had marched with the preceding VR session and provided ratings on measures of self-other blurring measured via the IOS Scale (49). The IOS Scale is a 7-point Scale with progressively overlapping circles to indicate degree of overlap between self and group presented. We also measured ratings of connection, likeability, similarity in personality, interest, marching synchrony feeling, and perceived difficulty of the marching task on a 5 point Likert-Scale.

### 1.6 Procedure

Upon the participant’s arrival, a screening questionnaire was administered to assess their suitability for participation in a VR study, particularly regarding any history of motion sickness. Then they were provided with a consent form outlining their rights. Once participants were deemed suitable for VR, they provided their informed consent, and then received instructions on how to march and respond to the random dot motion trials. To ensure synchronisation between their movements and those of the virtually presented synchrony group, the experimenter demonstrated the appropriate marching technique (ceremonial style military marching) in accordance with a designated reference beat (60 bpm). Participants were explicitly instructed to base their decision-making solely on the movement of the dots. Subsequently, participants were fitted with the VR headset and the practice phase was initiated.

During the practice phase, participants initially found themselves alone in the virtual scene. Once the beat commenced, any deviations from the instructed marching technique, particularly in terms of arm and leg movements and synchronisation with the beat, were corrected. The practice phase comprised three parts. In the first part, participants marched alone for a duration of 45 seconds, followed by five random dot motion trials. This sequence was then repeated with the synchronous and then the asynchronous group. Consequently, the practice phase had 15 random dot motion trials, containing all difficulty levels.

After completing the practice block, the initial manipulation of synchrony started. Depending on the assigned counterbalanced design, either the synchronous or asynchronous group reappeared. Participants engaged in 45 seconds of marching, followed by 30 trials of the random dot motion task. This sequence was repeated five times, comprising a block. Each block took approximately 15 minutes to complete. Subsequently, participants had their VR headset removed, and were provided with a questionnaire to assess their experience using a desktop-computer. This questionnaire gathered feedback and allowed participants to have a seated break in between the VR phases.

Additionally, participants were given the option to take a 5-10 minute break before proceeding with the second block. After the break, participants put on the VR headset once again and continued with the second block, which followed the same procedure as the first block but involved a different group. The presentation of the synchronous and asynchronous group was blocked and counter-balanced.

### 1.7 Data Processing

Movement profiles were extracted from the two VR hand controllers held by the participant in each hand. These were run through a low-pass Butterworth filter with a cut-off frequency of 10 Hz. To identify the raised hand in each trial, we compared the maximum velocity of one arm with that of the other. If the maximum velocity of one arm exceeded the other by at least 50%, that arm was designated as the decision made on the trial. In cases where these criteria were not met, the trial was either coded as both hands raised or as missed, and subsequently excluded from the analysis. After extraction, each trial was individually inspected to ensure that there were no recording errors and to ascertain the accuracy of the derived decision. 2.96% of the trials had to be manually corrected. These corrections were mostly applied to trials following a marching block as participants took some time to come to rest, trials where both arms were raised, or trials with recording difficulties.

We determined the reaction time for movement onset by analysing the velocity and position profiles of the raised arm movement. To do this, we first calculated the maximum velocity and position reached during the trial. We then traced back to the point where the velocity dropped to 20% of its maximum and the position was at 30% of the maximum.After manually inspecting both the velocity and position profiles to verify the accuracy of the calculated reaction times, we excluded 2.7% of trials due to inaccuracy. Additionally, reaction times for manually coded decision trials were also eliminated. Based on pre-registered criteria, trials were excluded when both arms met the decision threshold criteria, when neither arm reached the decision threshold criteria and when trials were identified as outliers in reaction time using the interquartile range method.

## 2. Results

### 2.1 Conformity

The proportion of trials where participants conformed with the decision of the group were submitted to a 2 (synchrony) × 2 (task difficulty) × 2 (group correctness) repeated measures ANOVA. The results indicated a main effect for task difficulty, *F*(1,56) = 14.233, *p* < .001, η^*2*^ = 0.005, and group correctness, *F*(1,56) = 979.426, *p* < .001, η^*2*^ = 0.584, but no main effect for synchrony, *F*(1,56) = 1.356, *p* = .249, η^*2*^ = 0.0004. Crucially, there was a significant synchrony × task difficulty interaction, *F*(1,56) = 7.001, *p* = .011, η^*2*^ = 0.002. Further examination of the post-hoc tests revealed that there was no difference in easy trials between synchronous and asynchronous marching conditions, *t*(56) = 0.919, *p*_holm_ > .05, *d* = 0.103, 95% CI [−0.197, 0.403]; in contrast, there was a significant difference in hard trials between synchronous and asynchronous marching conditions, *t*(56) = −2.642, *p*_holm_ = .038, *d* = −0.296, 95% CI [−0.604, 0.012].

There was a significant three-way interaction between synchrony, group correctness, and block order (*F*_(1,56)_ = 5.23, *p* = 0.026). This means the effects of synchrony and correctness varied depending on the order in which participants experienced the blocks (synchronous or asynchronous).When participants were exposed to incorrect group decisions, they gradually stopped conforming to those decisions over time, regardless of whether they started with the synchronous or asynchronous block. However, when the group was correct, the order of blocks made a difference. If participants began with the synchronous block, their conformity remained stable, even when they later moved to the asynchronous block. In contrast, if they started with the asynchronous block, their conformity significantly increased when they transitioned to the synchronous block. This suggests that participants were more likely to follow correct group decisions after moving from an asynchronous to a synchronous setting.The stable conformity rates when starting with the synchronous block may indicate that experiencing group synchrony generally enhances the perceived credibility of the group. The increased conformity when switching from asynchronous to synchronous blocks suggests that participants might perceive the synchronous group as more competent, especially after having experienced the asynchronous group. However, post-hoc pairwise tests did not reveal any statistically significant differences.

Trails with no correct responses, i.e, ambiguous trials were analysed separately. Ambiguous trials were submitted to a t-test comparing synchronous versus asynchronous condition (t_(57)_ = −1.39, p > 0.05), even though not significant, the test revealed a trend towards slightly higher levels of conformity following synchronous marching (M = .55, SD = .1) compared to asynchronous marching (M = .57, SD = .09).

### 2.2 Reaction Time

Reaction times were submitted to a 2×2×2 repeated measures ANOVA. The results indicated a main effect for task difficulty (*F*_(1,56)_ = 76.88, *p* < .001) but no main effect for of group correctness (*F*_(1,56)_ = 2.75, *p* = .01) and synchrony (*F*_(1,56)_ = .199, *p* = .657). Indicating that participants provided slower responses when the task was more difficult.

There was a significant synchrony × task difficulty interaction (*F*_(1,56)_ = 9.305, *p* = .003), indicating that in easy trials participants in the synchronous marching condition answered faster but this difference dissipated on hard trials. Furthermore, participants showed a significant improvement in reaction time across both blocks, with a significant synchrony × block order interaction (F(1,56) = 48.541, p < .001). Those who began with synchronous marching responded faster in both blocks compared to participants who started with asynchronous marching.

Reaction times decreased for both groups as the experiment progressed, but synchronous marching provided a consistent advantage On ambiguous trials, there was no difference in reaction times following synchronous versus asynchronous marching (t_(57)_ = .035, p > 0.05).

### 2.3 Accuracy

Task accuracy was submitted to a 2×2×2 repeated measures ANOVA. The results indicated a main effect for task difficulty (*F*_(1,56)_ = 1084.9, *p* < .001) and group correctness (*F*_(1,56)_ = 22.24, *p* < .001) but no main effect for synchrony (*F*_(1,56)_ = 3.178, p > 0.05). Participants were more accurate when the task was easy and when the group was correct. There was a significant task difficulty × group correctness interaction (*F*_(1,56)_ = 14.233, *p* < .001), this suggests that participants’ accuracy did not differ much based on group correctness in the easy trials, possibly due to ceiling effects. However, in the hard trials, participants’ accuracy was significantly higher when the group was correct.

There was a significant three way interaction between synchrony × task difficulty × group correctness (*F*_(1,53)_ = 7.001, *p* = .011), indicating that on hard trials when the group was correct and synchronous - participant accuracy was higher than when the group was correct and asynchronous. When the group was incorrect and synchronous, participant accuracy was slightly lower than when the group was incorrect and asynchronous. When the task was easy, participants’ accuracy was comparable across synchronous and asynchronous conditions. Pairwise post-hoc tests did not reveal any notable significant contrasts.

Additionally, there was a significant interaction between synchrony × block order (*F*_(1,56)_ = 5.238, *p* = .026). Further analysis of pairwise post-hoc tests indicated that this effect was primarily driven by participants who began with the asynchronous marching condition; their accuracy improved in the second block when surrounded by the synchronous group. Specifically, for participants who started with asynchronous marching, there was a significant difference in accuracy between the first and second block (t_(56)_ = −2.879, p_holm_ = .034). In contrast, participants who started with synchronous marching did not exhibit this pattern of higher accuracy in the second block (t_(56)_ = .358, p_holm_ > 0.5). This suggests that the order of experiencing synchronous marching plays a crucial role in accuracy improvement, with participants benefiting more when transitioning from an asynchronous to a synchronous group, whereas those who started with synchronous marching did not experience the same gains in accuracy.

### 2.4 Inverse Efficiency Score

Inverse Efficiency scores are calculated by dividing the mean RT of correct responses with the proportion of correct responses. It is an integrated measure of performance, lower values imply better performance. IES was submitted to a 2×2×2 repeated measures ANOVA. The results indicated a main effect for task difficulty (*F*_(1,53)_ = 149.135, *p* < .001) and group correctness (*F*_(1,53)_ = 19.269, *p* < .001) but no main effect for synchrony (*F*_(1,53)_ = 0.576, *p* = 0.451). There was a significant task difficulty × group correctness interaction (*F*_(1,53)_ = 18.321, *p* < .001). There was significant synchrony × block order interaction (*F*_(1,53)_ = 19.944, *p* < .001), since the synchrony manipulation is blocked this indicates that participants’s performance became better with time. There was a significant three way interaction between synchrony × group correctness × block order (*F*_(1,53)_ = 7.851, *p* = .007). There was a significant four way interaction synchrony × task difficulty ×group correctness × block order (*F*_(1,53)_ = 13.736, *p* < .001)

### 2.5 Post-Task Questionnaire Ratings

IOS Scale ratings were compared between the Synchronous marching and Asynchronous marching conditions. The distribution of IOS ratings was found to deviate from normality according to the Shapiro-Wilk test (W = 0.943, p = .009). Therefore, a paired Wilcoxon signed-rank test was conducted to compare the IOS ratings for the synchronous marching group (M = 2.93, SD = 1.29) and the asynchronous marching group (M = 2.47, SD = 1.273). The test revealed a significant difference between the two conditions (W = 590, p = .006, one-tailed, Sync > Async), indicating that the IOS ratings were significantly higher in the synchronous marching condition. See Fig. 3.

**Figure 3.**
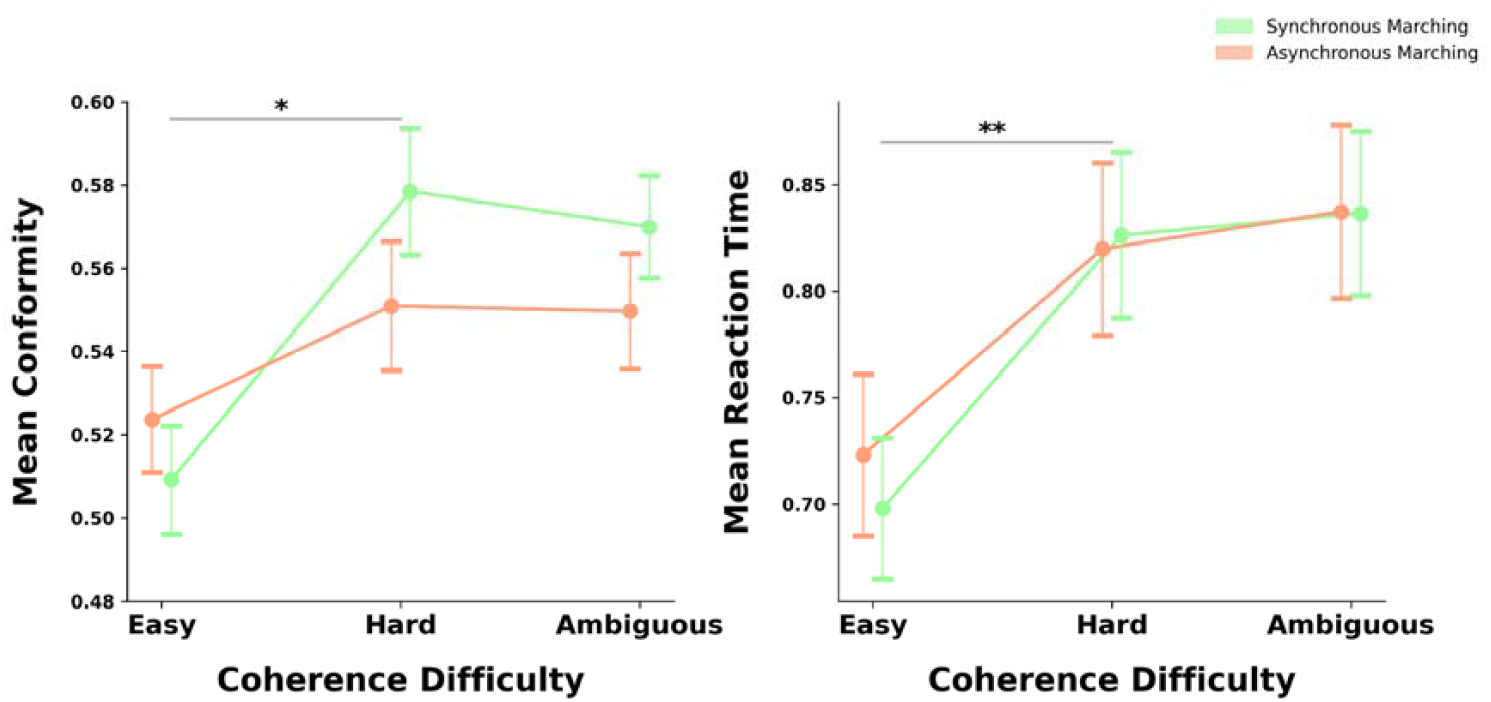
Left Panel: Mean Conformity across difficulty levels. Right Panel: Reaction Time. Error bars are standard errors f the mean (SEMs). * p < .05; ** p < .01.

**Figure 4.**
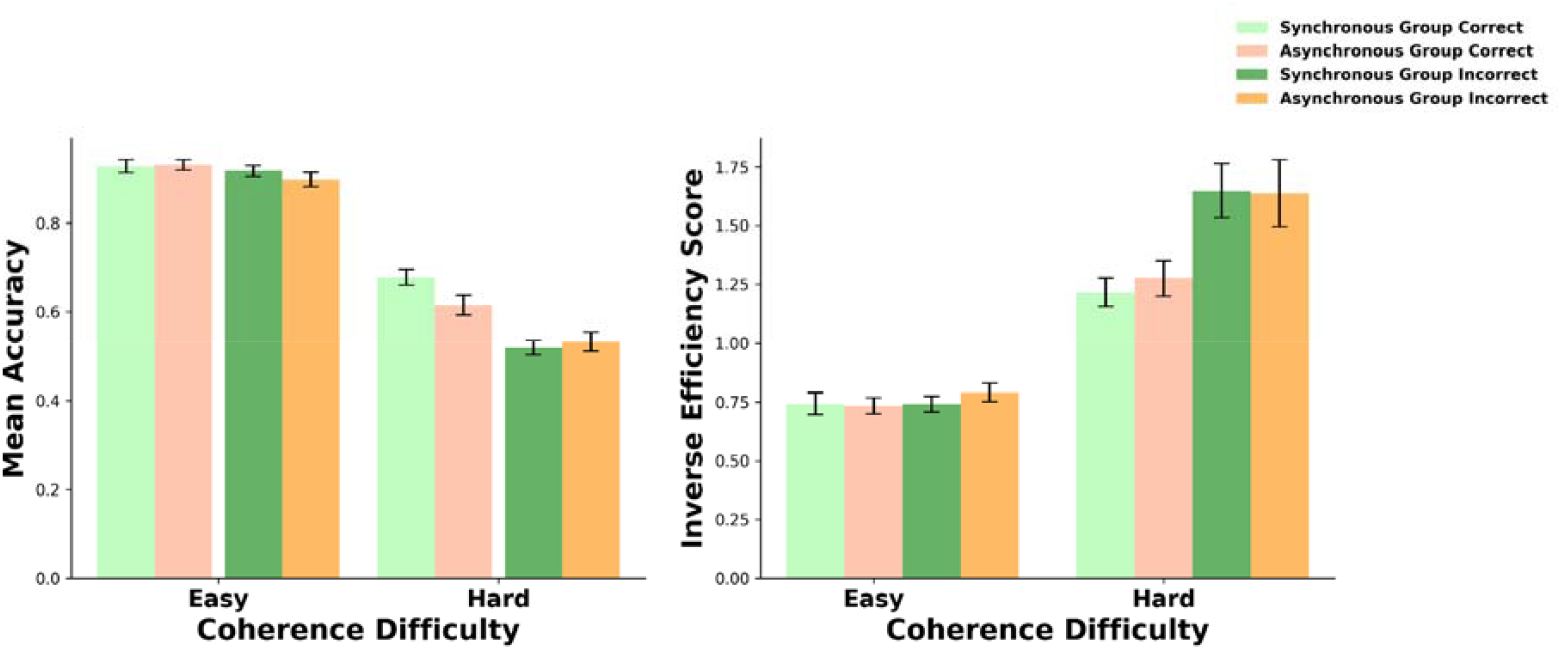
Left Panel: Mean Accuracy across difficulty and synchrony levels. Right Panel: Inverse Efficiency Scores cross difficulty and synchrony levels.. Error bars are standard errors of the mean (SEMs).

**Figure 4.**
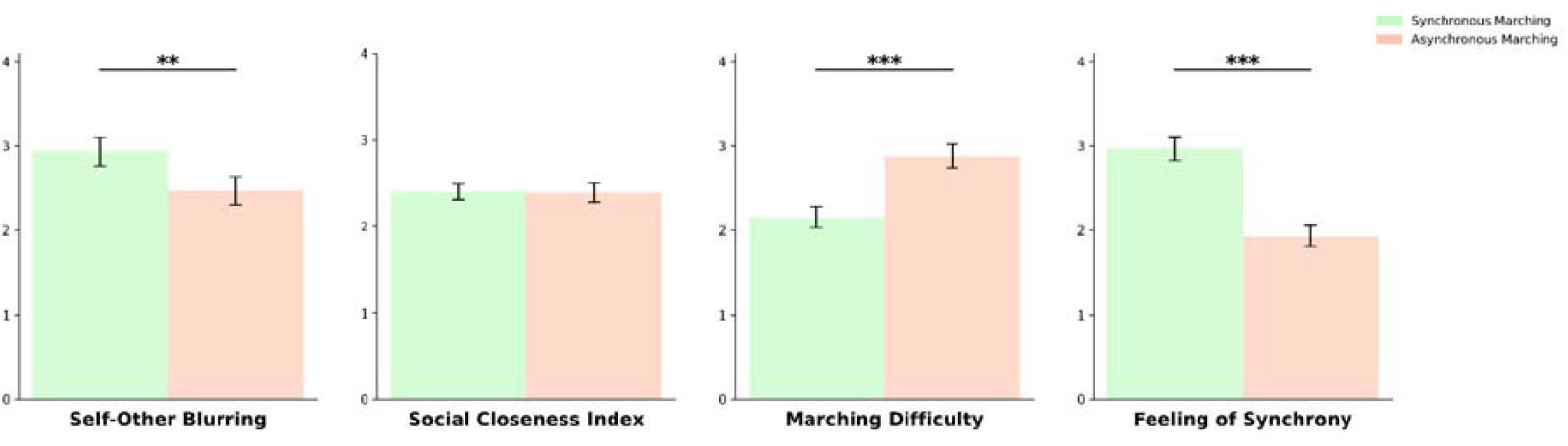
Self-Report Ratings. (1) Self-Other Blurring, (2) Social Closeness Index, (3) Marching difficulty, (4) Feeling of Synchrony while marching. Error bars are standard errors of the mean (SEMs).** p < .01; *** p < .001.

Ratings of connectedness, likability, similarity and interest were averaged to create the Social Closeness Index (SCI). We compared SCI between the Synchronous marching and Asynchronous marching conditions. A student-t test was conducted to compare the SCI scores for the synchronous marching group (M = 2.40, SD = .698) and the asynchronous marching group (M = 2.25, SD = .839). Results revealed no significant difference between synchronous and asynchronous conditions (t_(57)_= .131, p = .448, one-tailed Sync > Async).

Humans tend to naturally synchronise (50) making it difficult to maintain asynchronous behaviour. To assess the effectiveness of our experimental manipulation, participants rated their sense of synchrony and reported feeling more in sync with the group during synchronous marching. A comparison of Feeling of Synchrony ratings between the synchronous and asynchronous conditions showed that these ratings were not normally distributed, as indicated by the Shapiro-Wilk test (W = 0.914, p < .001). A paired Wilcoxon signed-rank test revealed a significant difference between the conditions (W = 848, p < .001, one-tailed Sync > Async).

Participants also rated asynchronous marching as more challenging than synchronous marching. The comparison of Marching Difficulty ratings between the two conditions similarly showed non-normal distribution (W = 0.905, p < .001), and the paired Wilcoxon signed-rank test confirmed a significant difference (W = 759, p < .001, one-tailed Sync < Async).

Thus, our analysis found significant differences in both the Feeling of Synchrony and Marching Difficulty between the synchronous and asynchronous conditions, which implies that participants considered the movements of the group during the marching phase while carrying out their own instructed behaviour.

## Discussion

Our results show that marching synchronously as compared to asynchronously increases conformity on an unrelated perceptual task under conditions of low stimulus information. This suggests that synchronised movement heightens susceptibility to group decision influence, adding to evidence that early perceptual faculties can be shaped by group identification (34,51). This study demonstrates the far-reaching consequences of collective movement-based group rituals. While it had been previously demonstrated that large-scale group rituals play a pivotal role in forging cohesive groups and reshaping personal identity (52), we have demonstrated that these rituals not only influence personal identity but also mould individual perception. Rituals that incorporate interpersonal movement synchrony and facilitate group induction through self-categorisation create unity and connection. The heightened sense of connection experienced during movement-based rituals inculcates greater trust in group decisions, establishing an optimal environment for inculcating a shared worldview. When people feel a stronger connection to a group, it creates a shared context and amplifies the significance of group norms, prompting individuals to align their judgments with the perceived consensus (53,54). In our evolutionary past, movement-based rituals likely functioned as an adaptive mechanism, part of our constitutional sociality, promoting the formation of highly cohesive groups and improving our chances of survival (10). However, in contemporary times, these ritual experiences may skew our perception by instilling a tendency to view the world through *‘group-coloured glasses’*

(51). This explains how messages from in-group members are more readily believed and better recalled (54), potentially contributing to the acceptance of false beliefs (55). We have demonstrated that even brief participation in movement-based rituals can impact perception, this suggests that with prolonged exposure, individuals may increasingly align their perceptions with group opinions, potentially prioritising these over objective reality, as demonstrated by studies such as Ross & Levy (2023), where participants adopted group norms in their judgments of reality.

Self-other blurring reflects a higher level of self-categorization within a synchronised group, which is a key factor in the group’s social influence. In the present experiment, participants reported greater self-other blurring when interacting with synchronised partners, aligning with previous research on interpersonal synchrony. This finding suggests that our manipulation taps into the same underlying mechanisms identified in the broader literature, where synchrony is frequently linked to a blurring of self-other boundaries (for review: Cross et al., 2019; Mogan et al., 2017). Moreover, our findings add to growing evidence that the influence of group identity on perceptual processes extends beyond long-standing groups, as even temporary or minimal groups created in experimental settings can distort perceptual decisions by fostering a sense of belonging and identification (51,57). While perceptual decision-making is often viewed as primarily driven by basic sensory inputs, our study suggests it is not immune to complex cognitive processes like group identity. The flexibility of our perceptual systems makes us vulnerable to social influence (46,58), which explains why group membership can affect how we perceive objective realities. Thus, experimentally induced synchronous activity serves as a group induction mechanism capable of distorting perceptual judgments. Of particular relevance to our study are paradigms that have demonstrated that minimal group identification causes perceptual distortions, specifically, a readiness to conform with the perceptual decisions of a minimally identified group (51). While continual participation in synchrony-based group rituals may create long lasting affiliations with groups (10), it can also trigger self-categorisation into experimentally induced artificial groups, or minimal groups (for review: Cross et al., 2019). Self-categorisation into minimal groups allows researchers to isolate the specific influence of group membership, disentangling it from factors like familiarity, experience, and expertise (59,60). According to the minimal group paradigm, cues are employed by individuals to categorise others even when group assortment is unrelated to the task at hand (61). Leading to expressive behaviours of in-group favouritism like increased resource allocation, trustworthiness and discrimination against the out-group (62).

We found no significant differences in social closeness across the synchrony conditions, a surprising result given that prior research has linked interpersonal synchrony with heightened feelings of social closeness (37). One possible explanation for this discrepancy could be that participants had pre-existing beliefs about the marching activity used in the experiment. Marching is often associated with military settings, which may have influenced how participants felt about the group, particularly in terms of similarity and likability. This strong association might have affected their responses, overshadowing any potential impact of synchrony. To avoid this in future studies, researchers should consider using activities that don’t carry such strong, specific connotations or measure participants’ beliefs about the activity beforehand to account for these biases.

In our experiment, both groups were deliberately programmed to be unreliable information sources with an accuracy of 50%. Thus, the rational choice for participants would be to disregard the group’s decisions due to their unreliability. However, our findings indicate that group correctness did influence participant decisions, particularly when synchrony was involved. When participants marched synchronously with a correct group, their own accuracy increased compared to when the group was asynchronous. Conversely, if the synchronous group was incorrect, participants’ accuracy decreased more than when the incorrect group was asynchronous. This suggests that synchrony amplifies the group’s influence on individual decision-making, potentially overriding rational considerations to disregard unreliable sources. Furthermore, the sequence in which participants experienced synchronous and asynchronous marching significantly impacted their task accuracy. Participants who began with asynchronous marching showed a notable improvement in accuracy during the second block when they transitioned to synchronous marching. In contrast, those who started with synchronous marching did not exhibit similar gains.

This indicates that moving from asynchrony to synchrony enhances sensitivity to group dynamics and the effectiveness of social cues on performance. This highlights the powerful role of synchrony in shaping social influence, even when it may not be rational to follow the group.

Future experimental designs could benefit from investigating the influence of synchrony on decision-making in the presence of reliable partners by enhancing group accuracy on easy trials. We programmed the groups in our experiment to show chance performance and nevertheless found an influence of the synchronous group. This strategic choice aimed to rigorously test our primary variable of interest, synchronous marching. Thus, our experimental manipulation specifically focuses on exploring how synchrony impacts the social influence exerted by unreliable partners. The high accuracy observed on easy trials suggests that the task was straightforward, and group inaccuracies during these trials could lead participants to disregard group decisions over time. Another intriguing avenue for exploration could comprise continuously adjusting task difficulty to identify where the shift in group reliance occurs and what role self-categorisation plays in this dynamic. Broadening the scope of our inquiry, additional investigations could systematically vary factors such as group size, diverse movement artefacts, timing of individual responses, and the diversity of opinions within a group (63). Furthermore, incorporating elements highlighted in previous research on social influence, such as participants’ beliefs about the nature of group decisions, whether automated or human (64) and the impact of social exclusion (65), would enhance the ecological validity of our paradigm, providing a comprehensive understanding of how group dynamics shape social influence. While investigating the separate effects of rhythm and movement artefacts could be a potential focus for future research, it falls outside the current study’s scope of interest

## Conclusion

In conclusion, our experiment demonstrates that participation in ritual experiences involving synchronous movement can shift perceptual decision-making to align more closely with synchronised partners. Interpersonal synchrony serves as a similarity cue that triggers self-categorisation into groups with synchronised partners, thereby enhancing the salience of group norms and making synchronised partners appear more credible. These findings provide insight into how ritual experiences contribute to groupthink and have broad implications beyond the laboratory, highlighting the far-reaching consequences of participating in group rituals. Understanding these dynamics is essential, particularly when considering tightly knit groups such as military organisations or cults, where informational conformity may drive radicalisation. The rapid adoption of group beliefs may explain events like riots and demonstrations, where collective synchronised actions like chanting or salutes may deepen group identity and render individuals more pliable to accept the reality presented by the group, potentially distorting perceptions and leading to collective rule-breaking. Conversely, this knowledge can be applied positively by incorporating synchronised movement in team-building activities or educational settings to enhance group cohesion (66,67).

## Acknowledgements and Funding Information

MBi was supported by the German Academic Exchange Service (DAAD; German: Deutscher Akademischer Austauschdienst). Additionally, this study was funded by DAAD, German Research Foundation (Deutsche Forschungsgemeinschaft, DFG) under Germany’s Excellence Strategy-EXC 2002/1, Science of Intelligence (Project Ref.: 390523135) and the Einstein Foundation Berlin. Further thanks to Valerii Chirkov (experiment development) and Dr.

Michael Galang (protocol advice). The funders had no role in study design, data collection and analysis, decision to publish, or preparation of the manuscript.

## Declarations

The authors have no competing interests to declare that are relevant to the content of this article.

## Author contributions

M.Bi.: Conceptualization, Writing – Original Draft, Writing – Review & Editing. M.Br.: Conceptualization, Writing – Review & Editing, Supervision, Funding acquisition.

