## SupplementaryMaterials1 for "Shared Rhythm to Shared Vision: Synchronous Marching increases Conformity on Perceptual Decision making"

**Supplementary Information 1**

**Section 1: Results**

**1.1 Accuracy**

Task accuracy was submitted to a 2×2×2 repeated measures ANOVA. The results indicated a main effect for task difficulty, *F*(1,56) = 1084.927, *p* < .001, *η²* = 0.595, and group correctness, *F*(1,56) = 22.240, *p* < .001, *η²* = 0.027, but no main effect for synchrony, *F*(1,56) = 3.178, *p* = .080, *η²* = 0.002, indicating that participants were more accurate when the task was easy and when the group was correct. There was a significant task difficulty × group correctness interaction, *F*(1,56) = 14.233, *p* < .001, *η²* = 0.012. Participants' accuracy did not differ much based on group correctness on the easy trials, possibly due to ceiling effects (*t*(56) = -2.113, *p*_holm_ = .111, *d* = -0.270, 95% CI [-0.617, 0.077]); however, on the hard trials, participants' accuracy was significantly higher when the group was correct (*t*(56) = -4.603, *p*_holm_ < .001, *d* = -0.566, 95% CI [-0.924, -0.209]).

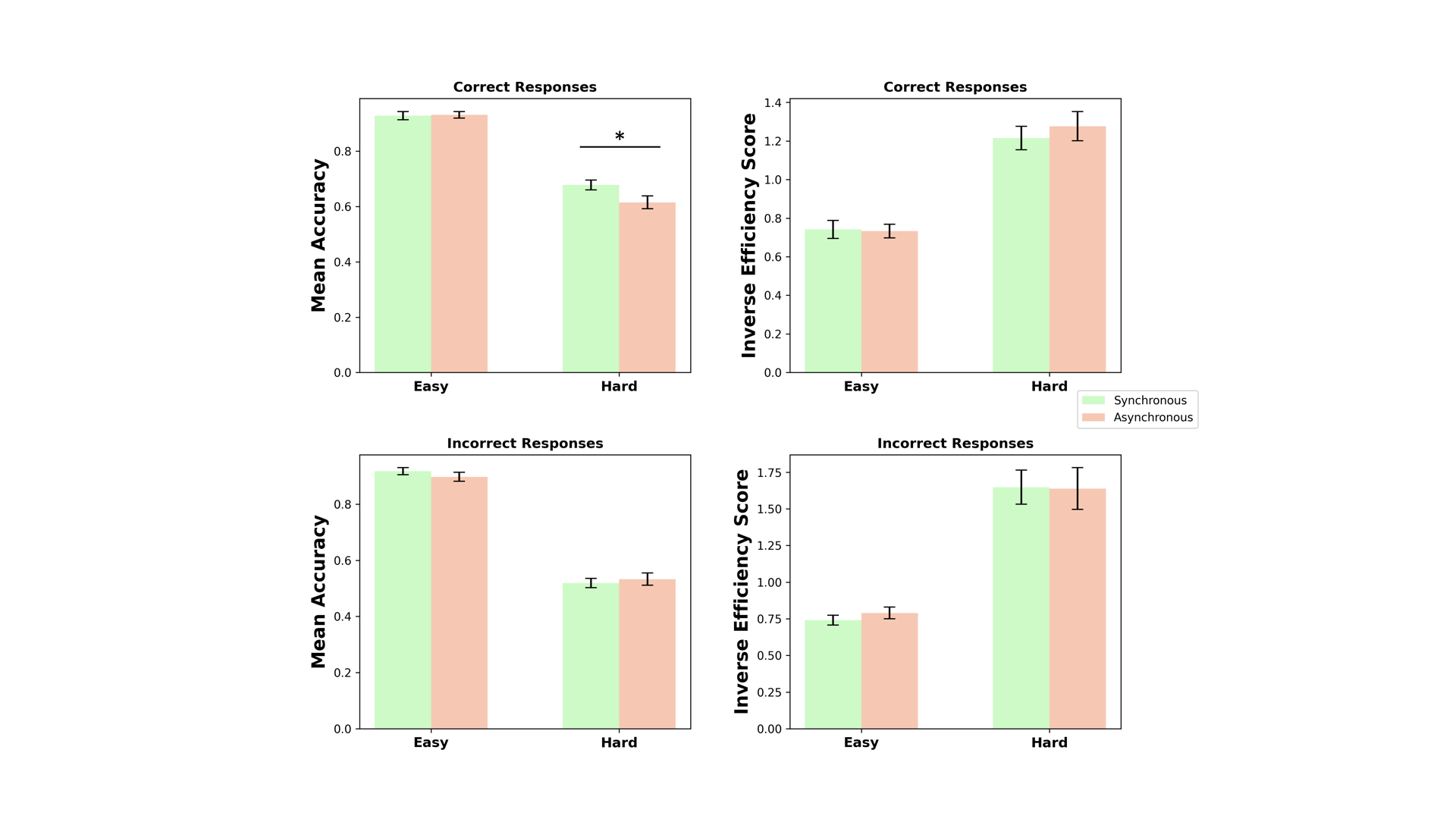

**Figure S1.** Left Panel: Mean Accuracy across difficulty and synchrony levels. Right Panel: Inverse Efficiency Scores across difficulty and synchrony levels.. Error bars are standard errors of the mean (SEMs).

There was also a significant three-way interaction between synchrony, task difficulty, and group correctness, *F*(1,56) = 7.001, *p* = .011, *η²* = 0.004. In hard trials, participant accuracy was significantly higher when the group was correct and synchronous (M = 0.67, SD = 0.13) compared to the asynchronous condition (M = 0.61, SD = 0.17), t(56) = -3.12, p_holm_ = .016, d = -0.49, 95% CI [-1.00, 0.03]. Other pairwise post-hoc tests did not reveal any notable significant contrasts (see Fig. S1).

Additionally, there was a significant interaction between synchrony and block order, *F*(1,56) = 5.238, *p* = .026, *η²* = 0.003. Further analysis of pairwise post-hoc tests indicated that this effect was primarily driven by participants who began with the asynchronous marching condition; their accuracy significantly improved in the second block when they transitioned to the synchronous condition *t*(56) = -2.88, *p*_holm_ = .034, *d* = -0.31, 95% CI [-0.61, -0.01]. In contrast, participants who started with synchronous marching exhibited consistently high accuracy across both blocks, showing no significant change, *t*(56) = 0.36, *p*_holm_ > .50, *d* = 0.04, 95% CI [-0.25, 0.33]. This suggests that synchronous marching provided a stable accuracy advantage, while those who initially experienced asynchronous marching benefited from synchrony only after transitioning in the second block, ultimately reaching the same accuracy levels as the synchronous-first group (see Figure S2).

**1.2 Inverse Efficiency Score**

Inverse Efficiency Scores (IES) were calculated by dividing the mean reaction time (RT) of correct responses by the proportion of correct responses. This integrated measure of performance reflects efficiency, with lower values indicating better performance. IES was submitted to a 2×2×2 repeated measures ANOVA. The results indicated a main effect of task difficulty, *F*(1,53) = 142.926, *p* < .001, *η²* = 0.255, and a main effect of group correctness, *F*(1,53) = 18.215, *p* < .001, *η²* = 0.022. However, there was no significant main effect of synchrony, *F*(1,53) = 0.398, *p* = .531, *η²* = 0.0005.

There was a significant task difficulty × group correctness interaction, *F*(1,53) = 15.574, *p* < .001, *η²* = 0.015, suggesting that the effect of task difficulty on IES depended on whether the group provided correct responses. In the easy condition, group correctness did not significantly impact efficiency (*t*(53) = 0.526, *p*_holm_ = 0.600, *d* = 0.060, 95% CI [-0.244, 0.364]). However, in the hard condition, participants were significantly more efficient when the group was correct compared to when it was incorrect (*t*(53) = 5.812, *p*_holm_ < .001, *d* = 0.659, 95% CI [0.310, 1.008]). Additionally, there was a significant synchrony × group correctness interaction, *F*(1,53) = 6.196, *p* = .016, *η²* = 0.003 and, a three-way interaction between synchrony, task difficulty, and group correctness was observed, *F*(1,53) = 11.137, *p* = .002, *η²* = 0.005, but post-hoc pairwise comparisons did not reveal any notable significant differences.

There was a significant synchrony × block order interaction, *F*(1,53) = 19.751, *p* < .001, *η²* = 0.026. Since the synchrony manipulation was blocked, this indicates that participants' performance improved over time. Post-hoc tests revealed that performance improved regardless of the direction of change, with participants who transitioned from asynchronous to synchronous marching showing significant improvement (*t*(53) = 3.494, *p*_holm_ = .006, *d* = 0.444, 95% CI [0.081, 0.808]), and those who transitioned from synchronous to asynchronous marching also demonstrating better performance compared to their first block (*t*(53) = -2.773, *p*_holm_ = .038, *d* = -0.334, 95% CI [-0.672, 0.004]). No other pairwise comparisons reached significance after correction. These findings suggest that the observed improvements in performance were not specific to the synchrony condition but instead reflect a general practice effect, with participants becoming better at the task over time.

There was a significant three-way interaction between synchrony, group correctness, and block order, *F*(1,53) = 6.196, *p* = .016, *η²* = 0.003. Additionally, a significant four-way interaction was found between synchrony, task difficulty, group correctness, and block order, *F*(1,53) = 11.137, *p* = .002, *η²* = 0.005. However, these higher-order interactions were difficult to interpret. Since synchrony was blocked, it completely overlapped with block order, making it difficult to determine whether these effects were driven by synchrony itself or simply a result of the block structure. Tables of the post-hoc tests can be found in the electronic supplementary information 2.

**1.3 Block order interaction plots**

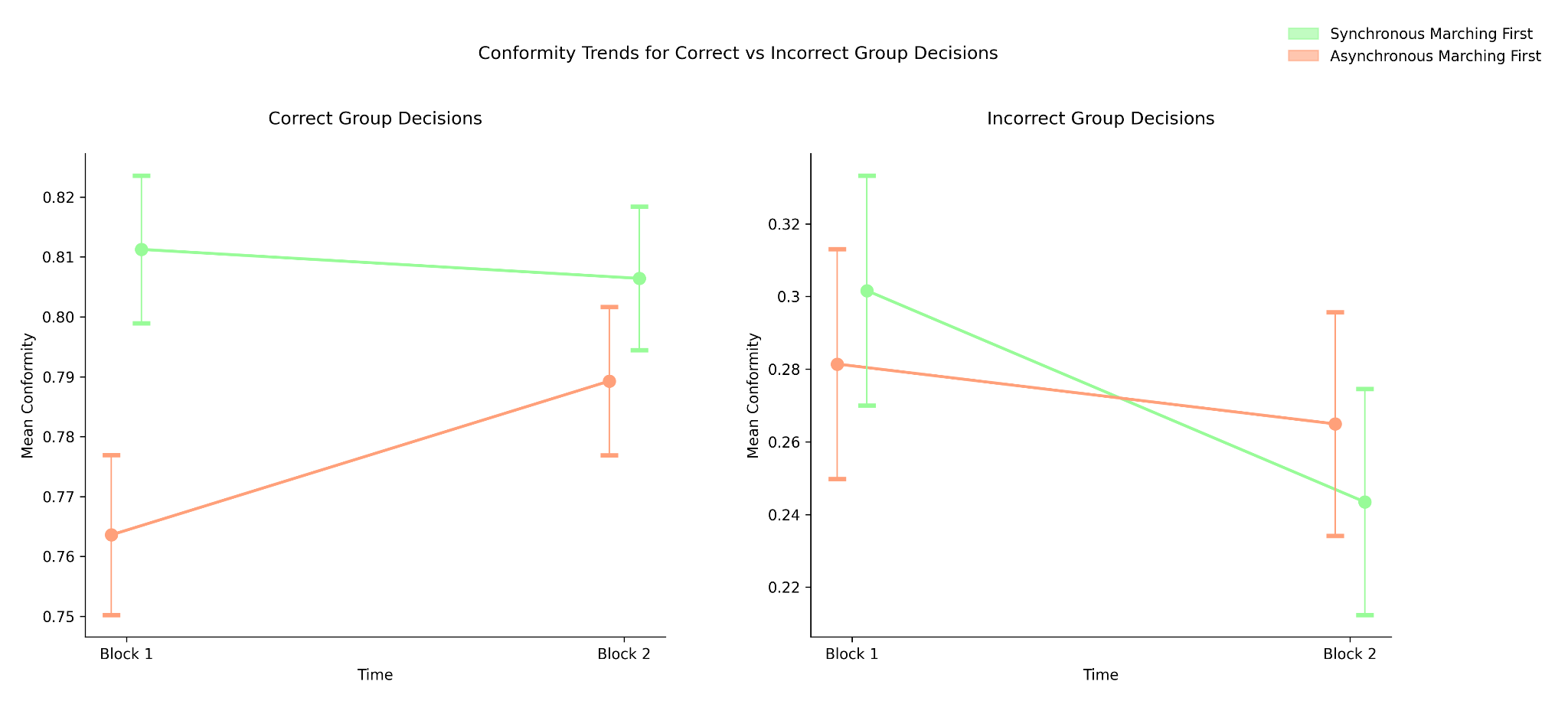

**Figure S2.** Mean Conformity Interaction Plot illustrating the synchrony × block × group correctness interaction. Left Panel: Correct Group Decisions; Right Panel: Incorrect Group Decisions. Error Bars represent standard error.

| 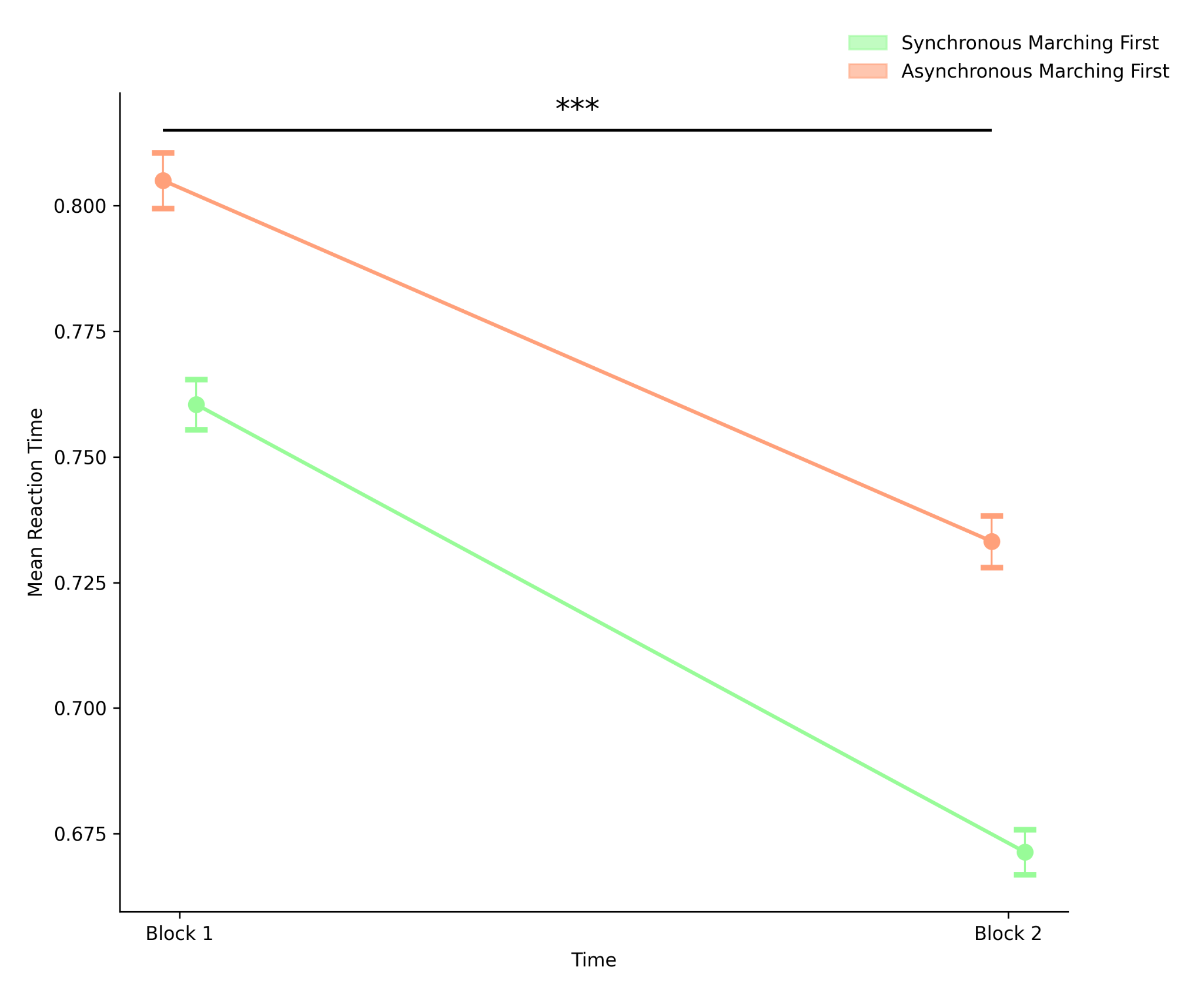 | 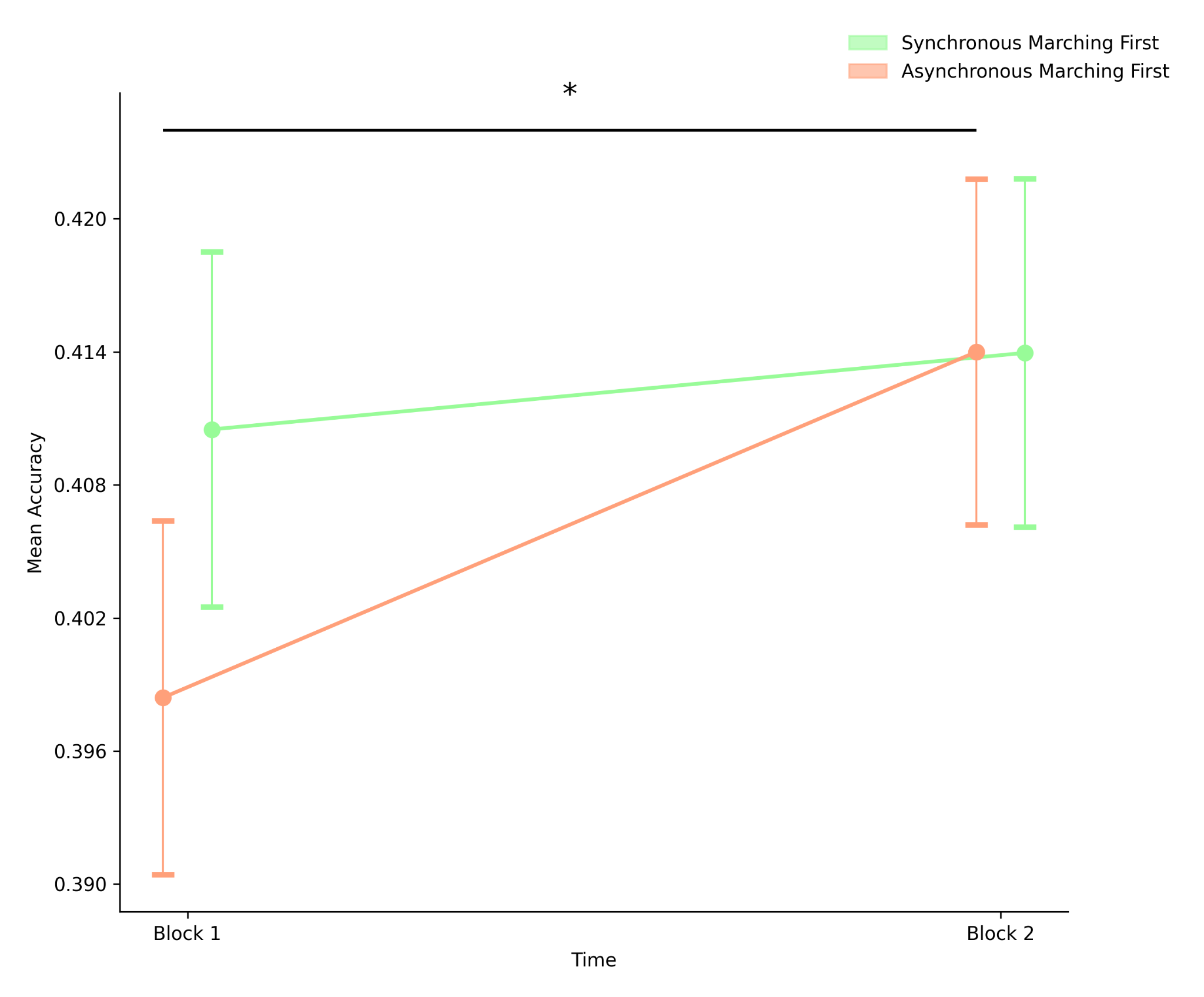 |
| --- | --- |
| **Figure S3.** Left Panel: Mean Reaction-time interaction plot illustrating the synchrony × block interaction; Right Panel: Mean Accuracy interaction plot illustrating the synchrony × block interaction. Error Bars represent standard error. | |

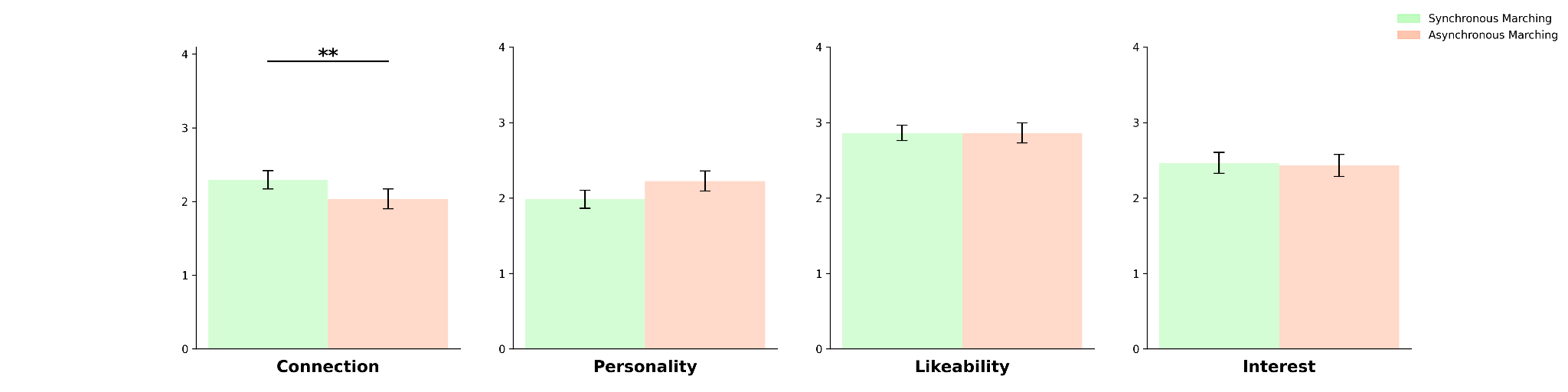

**Figure S4.** Likert Ratings for the composite social closeness index. (1) Connection, (2) Similar Personality, (3) Likeability, (4) Interest . Error bars are standard errors of the mean (SEMs).** p < .01

**1.4 Social Closeness Index**

Connection ratings were compared between the synchronous marching and asynchronous marching conditions. The distribution of connection ratings was found to deviate from normality based on the Shapiro-Wilk test (W = 0.924, p = .001). Results from a paired Wilcoxon signed-rank test to compare the connection ratings for the synchronous marching group (M = 2.29, SD = 0.95) and the asynchronous marching group (M = 2.03, SD = 1.04) revealed a significant difference between the two conditions (W = 487, p = 0.036, one-tailed, Sync > Async), indicating that the connection ratings were significantly higher in the synchronous marching condition. See Fig. S4.

Likeability ratings were compared between the synchronous marching and asynchronous marching conditions. The distribution of connection ratings was found to deviate from normality based on the Shapiro-Wilk test (W = 0.83, p < .001). Results from a paired Wilcoxon signed-rank test to compare the likeability ratings for the synchronous marching group (M = 2.86, SD = .78) and the asynchronous marching group (M = 2.86, SD = 1.03) revealed no significant difference between the two conditions (W = 137, p = 0.519, one-tailed, Sync > Async), indicating that the likeability ratings were not higher in the synchronous marching condition. See Fig. S4.

Ratings of similarity in terms of personality ratings were compared between the synchronous marching and asynchronous marching conditions. The distribution of connection ratings was found to deviate from normality based on the Shapiro-Wilk test (W = 0.927, p = .002). Results from a paired Wilcoxon signed-rank test was conducted to compare the connection ratings for the synchronous marching group (M = 1.98, SD = .9) and the asynchronous marching group (M = 2.22, SD = 1.02) revealed a no difference between the two conditions (W = 266, p = .94, one-tailed, Sync > Async), indicating that the ratings of similarity were not higher in the synchronous marching condition.

Interest ratings were compared between the synchronous marching and asynchronous marching conditions. The Shapiro-Wilk test indicated that the Interest scores were not normally distributed (W = 0.889, p < .001). Results from a paired Wilcoxon signed-rank test revealed no significant difference between the two conditions (W = 213, p = .416, one-tailed Sync > Async).

### **2. Sensitivity Analysis**

A sensitivity analysis was conducted using G*Power (Faul et al., 2007) to determine the minimum detectable effect size for the paired-samples t-test given α = 0.05, power = 0.80, and a sample size of 57. The analysis indicated that the study was sufficiently powered to detect an effect size of dz = 0.38. However, the observed effect size (dz = -0.296) was smaller than this threshold, suggesting that the study may have been underpowered to reliably detect effects of this magnitude. Given the novelty of this paradigm, future research should consider increasing statistical power, either by increasing sample size or by refining the experimental design to reduce variability and enhance sensitivity to smaller effects.

**3. Generalised Mixed Models**

Analysis was conducted on R.

**3.1 Conformity**

To identify the best-fitting model predicting GroupFollowing, we employed a generalised linear mixed model (GLMM) using the MuMIn package dredge() function for model selection. This approach systematically compared all possible models to determine the one with the best fit based on Akaike Information Criterion (AIC) and Bayesian Information Criterion (BIC) values. The models were fitted using the glmer() function from the lme4 package, with the binomial family (logit link function) and SubjectNumber as a random effect.

The structure of the best fitting model was:

*Conformity ~ CoherenceDifficulty + GroupCorrectness + synch +*

*(1 | SubjectNumber) + CoherenceDifficulty:GroupCorrectness + CoherenceDifficulty:synch*

Model Fit Statistics

| Model | AIC | BIC | logLik | Deviance | df |
| --- | --- | --- | --- | --- | --- |
| Final Model | 7883.2 | 7932.5 | -3934.6 | 7869.2 | 8358 |

| Predictor | Estimate | Std. Error | z-value | p-value |
| --- | --- | --- | --- | --- |
| (Intercept) | -2.1333 | 0.0951 | -22.423 | <0.001*** |
| CoherenceDifficulty (hard) | 1.9827 | 0.1052 | 18.841 | <0.001*** |
| GroupCorrectness (1) | 4.8953 | 0.1139 | 42.973 | <0.001*** |
| synch (TRUE) | -0.2263 | 0.1119 | -2.023 | 0.043* |
| CoherenceDifficulty × GroupCorrectness | -4.1743 | 0.1308 | -31.912 | <0.001*** |
| CoherenceDifficulty × synch | 0.3134 | 0.1291 | 2.427 | 0.015* |

Self-Report Variable Testing

After selecting the best model, we tested whether additional variables MarchSyncFeel, IOS, and SCI improved model fit. These variables were added individually and compared to the final model using ANOVA, AIC, and BIC values. The best model included CoherenceDifficulty, GroupCorrectness, and Synch, as well as the interaction terms CoherenceDifficulty × GroupCorrectness and CoherenceDifficulty × Synch. Attempts to improve model fit by adding MarchSyncFeel, IOS, and SCI did not yield significant improvements. Therefore, the final model remains the most parsimonious and best-fitting based on AIC/BIC values and statistical significance.

| Added Variable | AIC (Δ) | BIC (Δ) | LRT (p value) | Interpretation |
| --- | --- | --- | --- | --- |
| None (Best Model) | 7836.3 | 7871.4 | - | Baseline model |
| Perceived Synchrony | 7836.7 (+0.4) | 7878.9 (+7.5) | 0.2071 | No improvement |
| IOS | 7838.0 (+1.7) | 7880.2 (+8.8) | 0.6412 | No improvement |
| SCI | 7838.3 (+2.0) | 7880.4 (+9.0) | 0.9917 | No improvement |

**3.2 Conformity in Ambiguous trials**

In the analysis of ambiguous trials, we used a generalised linear mixed model (GLMM) to assess the effect of synchrony on group following. The model included synchrony as a fixed effect and SubjectNumber as a random effect. The results showed that the model had an AIC of 9717.7 and a BIC of 9738.3. The fixed effect of synchrony was not statistically significant (β = 0.06694, SE = 0.04816, z = 1.390, p = 0.165), indicating that synchrony did not significantly predict group following in ambiguous trials. Additionally, we tested whether adding Perceived Synchrony (MarchSyncFeel), Inclusion of Other in Self (IOS), and the Composite Social Closeness Index (SCI) would improve model fit. However, these additions did not lead to significant improvements (all p-values > 0.05). Therefore, the simplest model including only synchrony remained the most parsimonious representation of the data.

**3.3 Accuracy**

To identify the best-fitting model predicting AvatarCorrectness, we used a generalised linear mixed model (GLMM) and performed model selection using the MuMIn package's dredge() function. This method systematically compared all possible models to determine the best fit based on Akaike Information Criterion (AIC) and Bayesian Information Criterion (BIC) values. The models were fitted using the glmer() function from the lme4 package, with a binomial family (logit link function) and SubjectNumber as a random effect.

The structure of the best fitting model was:

*Accuracy ~ CoherenceDifficulty + GroupCorrectness + synch + (1 | SubjectNumber)*

Model Fit Statistics

| Model | AIC | BIC | logLik | Deviance | df |
| --- | --- | --- | --- | --- | --- |
| Final Model | 7836.3 | 7871.4 | -3913.1 | 7826.3 | 8360 |

Fixed Effects Results

| Predictor | Estimate | Std. Error | z-value | p-value |
| --- | --- | --- | --- | --- |
| (Intercept) | 2.2089 | 0.0800 | 27.608 | <0.001*** |
| CoherenceDifficulty (hard) | -2.1097 | 0.0651 | -32.433 | <0.001*** |
| GroupCorrectness (1) | 0.4664 | 0.0565 | 8.249 | <0.001*** |
| synch (TRUE) | 0.0813 | 0.0562 | 1.446 | 0.148 |

The model revealed that CoherenceDifficulty (hard) had a strong negative effect on AvatarCorrectness, indicating that harder trials led to lower accuracy. GroupCorrectness had a significant positive effect, meaning participants were more likely to be correct when group correctness was high. Synchrony (synch) was not a significant predictor of accuracy (p = .148), suggesting that synchrony had little to no direct impact on AvatarCorrectness.

Self-Report Variable Testing

After selecting the best model, we tested whether additional variables—MarchSyncFeel, IOS, and SCI—improved model fit. These variables were added individually and compared to the final model using ANOVA, AIC, and BIC values.None of these additional predictors significantly improved model fit.

| Added Variable | AIC (Δ) | BIC (Δ) | LRT (p value) | Interpretation |
| --- | --- | --- | --- | --- |
| None (Best Model) | 7836.3 | 7871.4 | - | Baseline model |
| Perceived Synchrony | 7836.7 (+0.4) | 7878.9 (+7.5) | 0.2071 | No improvement |
| IOS | 7838.0 (+1.7) | 7880.2 (+8.8) | 0.6412 | No improvement |
| SCI | 7838.3 (+2.0) | 7880.4 (+9.0) | 0.9917 | No improvement |
