## SupplementaryMaterials2 for "Shared Rhythm to Shared Vision: Synchronous Marching increases Conformity on Perceptual Decision making"

**Supplementary Information 2**

a) **Conformity – PostHoc Pairwairse Comparisons**

**Table S1.** Summary of the post-hoc pairwise comparisons for conformity in synchrony and difficulty conditions. The table presents the mean difference, 95% confidence intervals (CI), standard error (SE), t-value, Cohen's d effect size, and adjusted p-values (pholm) for multiple comparisons.

| *Post Hoc Comparisons - Synchrony ✻ Difficulty* | | | | | | | | | | | | | | | | | | | | | |
| --- | --- | --- | --- | --- | --- | --- | --- | --- | --- | --- | --- | --- | --- | --- | --- | --- | --- | --- | --- | --- | --- |
|  | | | | | | 95% CI for Mean Difference | | | |  | | | | | | 95% CI for Cohen's d | | | |  | |
|  | |  | | Mean Difference | | Lower | | Upper | | SE | | t | | Cohen's d | | Lower | | Upper | | p_holm_ | |
| async, easy |  | sync, easy |  | 0.013 |  | -0.026 |  | 0.052 |  | 0.015 |  | 0.919 |  | 0.103 |  | -0.197 |  | 0.403 |  | 0.360 |  |
|  |  | async, hard |  | -0.022 |  | -0.065 |  | 0.021 |  | 0.016 |  | -1.357 |  | -0.167 |  | -0.497 |  | 0.164 |  | 0.356 |  |
|  |  | sync, hard |  | -0.060 |  | -0.105 |  | -0.016 |  | 0.017 |  | -3.625 |  | -0.463 |  | -0.823 |  | -0.103 |  | 0.002 | ** |
| sync, easy |  | async, hard |  | -0.035 |  | -0.080 |  | 0.010 |  | 0.017 |  | -2.113 |  | -0.270 |  | -0.617 |  | 0.077 |  | 0.111 |  |
|  |  | sync, hard |  | -0.073 |  | -0.116 |  | -0.031 |  | 0.016 |  | -4.603 |  | -0.566 |  | -0.924 |  | -0.209 |  | < .001 | *** |
| async, hard |  | sync, hard |  | -0.038 |  | -0.077 |  | 6.492×10^-4^ |  | 0.015 |  | -2.642 |  | -0.296 |  | -0.604 |  | 0.012 |  | 0.038 | * |
| * p < .05, ** p < .01, *** p < .001 | | | | | | | | | | | | | | | | | | | | | |
| Note.  P-value and confidence intervals adjusted for comparing a family of 6 estimates (confidence intervals corrected using the bonferroni method). | | | | | | | | | | | | | | | | | | | | | |
| Note.  Results are averaged over the levels of: Blockfirst, Group Correctness | | | | | | | | | | | | | | | | | | | | | |

**Table S2.** Summary of the post-hoc pairwise comparisons for conformity across block order (Blockfirst), synchrony, and group correctness. The table includes the mean difference, 95% CI, SE, t-value, Cohen’s d, and p-value adjustments for multiple comparisons.

| *Post Hoc Comparisons - Blockfirst ✻ Synchrony ✻ Group Correctness* | | | | | | | | | | | | | | | | | | | | | |
| --- | --- | --- | --- | --- | --- | --- | --- | --- | --- | --- | --- | --- | --- | --- | --- | --- | --- | --- | --- | --- | --- |
|  | | | | | | 95% CI for Mean Difference | | | |  | | | | | | 95% CI for Cohen's d | | | |  | |
|  | |  | | Mean Difference | | Lower | | Upper | | SE | | t | | Cohen's d | | Lower | | Upper | | p_holm_ | |
| AsyncFirst, async, group incorrect |  | SyncFirst, async, group incorrect |  | 0.023 |  | -0.060 |  | 0.107 |  | 0.026 |  | 0.882 |  | 0.180 |  | -0.468 |  | 0.827 |  | 1.000 |  |
|  |  | AsyncFirst, sync, group incorrect |  | 0.031 |  | -0.035 |  | 0.097 |  | 0.021 |  | 1.522 |  | 0.242 |  | -0.267 |  | 0.750 |  | 1.000 |  |
|  |  | SyncFirst, sync, group incorrect |  | 0.002 |  | -0.082 |  | 0.086 |  | 0.026 |  | 0.071 |  | 0.014 |  | -0.630 |  | 0.659 |  | 1.000 |  |
|  |  | AsyncFirst, async, group correct |  | -0.446 |  | -0.532 |  | -0.360 |  | 0.027 |  | -16.744 |  | -3.440 |  | -4.659 |  | -2.222 |  | < .001 | *** |
|  |  | SyncFirst, async, group correct |  | -0.502 |  | -0.586 |  | -0.419 |  | 0.026 |  | -19.031 |  | -3.874 |  | -5.201 |  | -2.547 |  | < .001 | *** |
|  |  | AsyncFirst, sync, group correct |  | -0.495 |  | -0.583 |  | -0.407 |  | 0.027 |  | -18.076 |  | -3.815 |  | -5.138 |  | -2.492 |  | < .001 | *** |
|  |  | SyncFirst, sync, group correct |  | -0.514 |  | -0.598 |  | -0.430 |  | 0.026 |  | -19.466 |  | -3.963 |  | -5.313 |  | -2.613 |  | < .001 | *** |
| SyncFirst, async, group incorrect |  | AsyncFirst, sync, group incorrect |  | 0.008 |  | -0.076 |  | 0.092 |  | 0.026 |  | 0.305 |  | 0.062 |  | -0.583 |  | 0.707 |  | 1.000 |  |
|  |  | SyncFirst, sync, group incorrect |  | -0.021 |  | -0.087 |  | 0.045 |  | 0.021 |  | -1.040 |  | -0.165 |  | -0.670 |  | 0.340 |  | 1.000 |  |
|  |  | AsyncFirst, async, group correct |  | -0.469 |  | -0.553 |  | -0.386 |  | 0.026 |  | -17.782 |  | -3.620 |  | -4.881 |  | -2.359 |  | < .001 | *** |
|  |  | SyncFirst, async, group correct |  | -0.526 |  | -0.611 |  | -0.440 |  | 0.027 |  | -19.730 |  | -4.054 |  | -5.431 |  | -2.677 |  | < .001 | *** |
|  |  | AsyncFirst, sync, group correct |  | -0.518 |  | -0.602 |  | -0.434 |  | 0.026 |  | -19.621 |  | -3.994 |  | -5.353 |  | -2.636 |  | < .001 | *** |
|  |  | SyncFirst, sync, group correct |  | -0.537 |  | -0.625 |  | -0.449 |  | 0.027 |  | -19.629 |  | -4.142 |  | -5.551 |  | -2.734 |  | < .001 | *** |
| AsyncFirst, sync, group incorrect |  | SyncFirst, sync, group incorrect |  | -0.029 |  | -0.113 |  | 0.054 |  | 0.026 |  | -1.116 |  | -0.227 |  | -0.876 |  | 0.421 |  | 1.000 |  |
|  |  | AsyncFirst, async, group correct |  | -0.478 |  | -0.565 |  | -0.390 |  | 0.027 |  | -17.447 |  | -3.682 |  | -4.971 |  | -2.393 |  | < .001 | *** |
|  |  | SyncFirst, async, group correct |  | -0.534 |  | -0.617 |  | -0.450 |  | 0.026 |  | -20.218 |  | -4.116 |  | -5.506 |  | -2.725 |  | < .001 | *** |
|  |  | AsyncFirst, sync, group correct |  | -0.526 |  | -0.612 |  | -0.440 |  | 0.027 |  | -19.742 |  | -4.056 |  | -5.434 |  | -2.679 |  | < .001 | *** |
|  |  | SyncFirst, sync, group correct |  | -0.545 |  | -0.629 |  | -0.462 |  | 0.026 |  | -20.653 |  | -4.204 |  | -5.619 |  | -2.790 |  | < .001 | *** |
| SyncFirst, sync, group incorrect |  | AsyncFirst, async, group correct |  | -0.448 |  | -0.532 |  | -0.364 |  | 0.026 |  | -16.971 |  | -3.455 |  | -4.674 |  | -2.236 |  | < .001 | *** |
|  |  | SyncFirst, async, group correct |  | -0.504 |  | -0.592 |  | -0.416 |  | 0.027 |  | -18.427 |  | -3.889 |  | -5.231 |  | -2.546 |  | < .001 | *** |
|  |  | AsyncFirst, sync, group correct |  | -0.497 |  | -0.580 |  | -0.413 |  | 0.026 |  | -18.810 |  | -3.829 |  | -5.144 |  | -2.514 |  | < .001 | *** |
|  |  | SyncFirst, sync, group correct |  | -0.516 |  | -0.602 |  | -0.430 |  | 0.027 |  | -19.358 |  | -3.977 |  | -5.334 |  | -2.621 |  | < .001 | *** |
| AsyncFirst, async, group correct |  | SyncFirst, async, group correct |  | -0.056 |  | -0.140 |  | 0.027 |  | 0.026 |  | -2.132 |  | -0.434 |  | -1.092 |  | 0.224 |  | 0.343 |  |
|  |  | AsyncFirst, sync, group correct |  | -0.049 |  | -0.114 |  | 0.017 |  | 0.021 |  | -2.358 |  | -0.374 |  | -0.890 |  | 0.141 |  | 0.222 |  |
|  |  | SyncFirst, sync, group correct |  | -0.068 |  | -0.151 |  | 0.016 |  | 0.026 |  | -2.567 |  | -0.523 |  | -1.186 |  | 0.141 |  | 0.132 |  |
| SyncFirst, async, group correct |  | AsyncFirst, sync, group correct |  | 0.008 |  | -0.076 |  | 0.091 |  | 0.026 |  | 0.293 |  | 0.060 |  | -0.586 |  | 0.705 |  | 1.000 |  |
|  |  | SyncFirst, sync, group correct |  | -0.011 |  | -0.077 |  | 0.054 |  | 0.021 |  | -0.558 |  | -0.089 |  | -0.592 |  | 0.415 |  | 1.000 |  |
| AsyncFirst, sync, group correct |  | SyncFirst, sync, group correct |  | -0.019 |  | -0.103 |  | 0.064 |  | 0.026 |  | -0.728 |  | -0.148 |  | -0.795 |  | 0.498 |  | 1.000 |  |
| *** p < .001 | | | | | | | | | | | | | | | | | | | | | |
| *Note.*  P-value and confidence intervals adjusted for comparing a family of 28 estimates (confidence intervals corrected using the bonferroni method). | | | | | | | | | | | | | | | | | | | | | |
| *Note.*  Results are averaged over the levels of: Difficulty | | | | | | | | | | | | | | | | | | | | | |

b) **Reaction Time – PostHoc Pairwairse Comparisons**

**Table S3.** Summary of the post-hoc pairwise comparisons for reaction time across synchrony and difficulty conditions. The table presents statistical comparisons including mean difference, 95% CI, SE, t-value, Cohen’s d, and corrected p-values.

| *Post Hoc Comparisons - Synchrony ✻ Difficulty* | | | | | | | | | | | | | | | | | | | | | |
| --- | --- | --- | --- | --- | --- | --- | --- | --- | --- | --- | --- | --- | --- | --- | --- | --- | --- | --- | --- | --- | --- |
|  | | | | | | 95% CI for Mean Difference | | | |  | | | | | | 95% CI for Cohen's d | | | |  | |
|  | |  | | Mean Difference | | Lower | | Upper | | SE | | t | | Cohen's d | | Lower | | Upper | | p_holm_ | |
| async, easy |  | sync, easy |  | 0.025 |  | -0.021 |  | 0.070 |  | 0.017 |  | 1.463 |  | 0.088 |  | -0.077 |  | 0.252 |  | 0.296 |  |
|  |  | async, hard |  | -0.093 |  | -0.131 |  | -0.056 |  | 0.014 |  | -6.723 |  | -0.333 |  | -0.493 |  | -0.174 |  | < .001 | *** |
|  |  | sync, hard |  | -0.104 |  | -0.158 |  | -0.050 |  | 0.020 |  | -5.139 |  | -0.371 |  | -0.589 |  | -0.153 |  | < .001 | *** |
| sync, easy |  | async, hard |  | -0.118 |  | -0.172 |  | -0.064 |  | 0.020 |  | -5.835 |  | -0.421 |  | -0.645 |  | -0.197 |  | < .001 | *** |
|  |  | sync, hard |  | -0.129 |  | -0.166 |  | -0.091 |  | 0.014 |  | -9.245 |  | -0.459 |  | -0.638 |  | -0.279 |  | < .001 | *** |
| async, hard |  | sync, hard |  | -0.010 |  | -0.056 |  | 0.035 |  | 0.017 |  | -0.624 |  | -0.037 |  | -0.201 |  | 0.126 |  | 0.534 |  |
| *** p < .001 | | | | | | | | | | | | | | | | | | | | | |
| *Note.*  P-value and confidence intervals adjusted for comparing a family of 6 estimates (confidence intervals corrected using the bonferroni method). | | | | | | | | | | | | | | | | | | | | | |
| *Note.*  Results are averaged over the levels of: Blockfirst, Group Correctness | | | | | | | | | | | | | | | | | | | | | |

| **Table S4.** Summary of the post-hoc pairwise comparisons for reaction time across synchrony and block order (Blockfirst),. The table presents statistical comparisons including mean difference, 95% CI, SE, t-value, Cohen’s d, and corrected p-values.  *Post Hoc Comparisons - Blockfirst ✻ Synchrony* | | | | | | | | | | | | | | | | | | | | | |
| --- | --- | --- | --- | --- | --- | --- | --- | --- | --- | --- | --- | --- | --- | --- | --- | --- | --- | --- | --- | --- | --- |
|  | | | | | | 95% CI for Mean Difference | | | |  | | | | | | 95% CI for Cohen's d | | | |  | |
|  | |  | | Mean Difference | | Lower | | Upper | | SE | | t | | Cohen's d | | Lower | | Upper | | p_holm_ | |
| AsyncFirst, async |  | SyncFirst, async |  | 0.167 |  | -0.028 |  | 0.362 |  | 0.072 |  | 2.330 |  | 0.595 |  | -0.118 |  | 1.307 |  | 0.092 |  |
|  |  | AsyncFirst, sync |  | 0.117 |  | 0.056 |  | 0.178 |  | 0.022 |  | 5.242 |  | 0.417 |  | 0.175 |  | 0.660 |  | < .001 | *** |
|  |  | SyncFirst, sync |  | 0.064 |  | -0.131 |  | 0.259 |  | 0.072 |  | 0.892 |  | 0.228 |  | -0.471 |  | 0.926 |  | 1.000 |  |
| SyncFirst, async |  | AsyncFirst, sync |  | -0.050 |  | -0.245 |  | 0.145 |  | 0.072 |  | -0.695 |  | -0.177 |  | -0.875 |  | 0.520 |  | 1.000 |  |
|  |  | SyncFirst, sync |  | -0.103 |  | -0.164 |  | -0.042 |  | 0.022 |  | -4.611 |  | -0.367 |  | -0.604 |  | -0.130 |  | < .001 | *** |
| AsyncFirst, sync |  | SyncFirst, sync |  | -0.053 |  | -0.248 |  | 0.142 |  | 0.072 |  | -0.744 |  | -0.190 |  | -0.887 |  | 0.508 |  | 1.000 |  |
| *** p < .001 | | | | | | | | | | | | | | | | | | | | | |
| *Note.*  P-value and confidence intervals adjusted for comparing a family of 6 estimates (confidence intervals corrected using the bonferroni method). | | | | | | | | | | | | | | | | | | | | | |
| *Note.*  Results are averaged over the levels of: Difficulty, Group Correctness | | | | | | | | | | | | | | | | | | | | | |

c) **Accuracy– PostHoc Pairwairse Comparisons**

**Table S5.** Summary of the post-hoc pairwise comparisons for accuracy across difficulty and group correctness conditions. The table includes mean differences, 95% CI, SE, t-values, Cohen's d, and adjusted p-values.

| *Post Hoc Comparisons - Difficulty ✻ Group Correctness* | | | | | | | | | | | | | | | | | | | | | |
| --- | --- | --- | --- | --- | --- | --- | --- | --- | --- | --- | --- | --- | --- | --- | --- | --- | --- | --- | --- | --- | --- |
|  | | | | | | 95% CI for Mean Difference | | | |  | | | | | | 95% CI for Cohen's d | | | |  | |
|  | |  | | Mean Difference | | Lower | | Upper | | SE | | t | | Cohen's d | | Lower | | Upper | | p_holm_ | |
| easy, group incorrect |  | hard, group incorrect |  | 0.380 |  | 0.336 |  | 0.423 |  | 0.016 |  | 23.529 |  | 2.928 |  | 2.120 |  | 3.737 |  | < .001 | *** |
|  |  | easy, group correct |  | -0.023 |  | -0.076 |  | 0.029 |  | 0.020 |  | -1.192 |  | -0.180 |  | -0.586 |  | 0.225 |  | 0.236 |  |
|  |  | hard, group correct |  | 0.261 |  | 0.213 |  | 0.310 |  | 0.018 |  | 14.429 |  | 2.015 |  | 1.386 |  | 2.644 |  | < .001 | *** |
| hard, group incorrect |  | easy, group correct |  | -0.403 |  | -0.452 |  | -0.354 |  | 0.018 |  | -22.262 |  | -3.109 |  | -3.975 |  | -2.242 |  | < .001 | *** |
|  |  | hard, group correct |  | -0.118 |  | -0.171 |  | -0.066 |  | 0.020 |  | -6.038 |  | -0.914 |  | -1.378 |  | -0.449 |  | < .001 | *** |
| easy, group correct |  | hard, group correct |  | 0.285 |  | 0.241 |  | 0.328 |  | 0.016 |  | 17.639 |  | 2.195 |  | 1.551 |  | 2.840 |  | < .001 | *** |
| *** p < .001 | | | | | | | | | | | | | | | | | | | | | |
| *Note.*  P-value and confidence intervals adjusted for comparing a family of 6 estimates (confidence intervals corrected using the bonferroni method). | | | | | | | | | | | | | | | | | | | | | |
| *Note.*  Results are averaged over the levels of: Blockfirst, Synchrony | | | | | | | | | | | | | | | | | | | | | |

**Table S6.** Summary of the post-hoc pairwise comparisons for accuracy across synchrony, difficulty, and group correctness conditions. The table details mean differences, 95% CI, SE, t-values, Cohen’s d, and p-value corrections.

| *Post Hoc Comparisons - Synchrony ✻ Difficulty ✻ Group Correctness* | | | | | | | | | | | | | | | | | | | | | |
| --- | --- | --- | --- | --- | --- | --- | --- | --- | --- | --- | --- | --- | --- | --- | --- | --- | --- | --- | --- | --- | --- |
|  | | | | | | 95% CI for Mean Difference | | | |  | | | | | | 95% CI for Cohen's d | | | |  | |
|  | |  | | Mean Difference | | Lower | | Upper | | SE | | t | | Cohen's d | | Lower | | Upper | | p_holm_ | |
| async, easy, group incorrect |  | sync, easy, group incorrect |  | -0.023 |  | -0.088 |  | 0.041 |  | 0.020 |  | -1.143 |  | -0.179 |  | -0.676 |  | 0.317 |  | 1.000 |  |
|  |  | async, hard, group incorrect |  | 0.361 |  | 0.293 |  | 0.430 |  | 0.022 |  | 16.801 |  | 2.787 |  | 1.808 |  | 3.767 |  | < .001 | *** |
|  |  | sync, hard, group incorrect |  | 0.375 |  | 0.306 |  | 0.444 |  | 0.022 |  | 17.243 |  | 2.890 |  | 1.882 |  | 3.898 |  | < .001 | *** |
|  |  | async, easy, group correct |  | -0.037 |  | -0.114 |  | 0.041 |  | 0.024 |  | -1.505 |  | -0.283 |  | -0.882 |  | 0.315 |  | 0.937 |  |
|  |  | sync, easy, group correct |  | -0.033 |  | -0.110 |  | 0.043 |  | 0.024 |  | -1.373 |  | -0.257 |  | -0.850 |  | 0.336 |  | 1.000 |  |
|  |  | async, hard, group correct |  | 0.281 |  | 0.207 |  | 0.356 |  | 0.023 |  | 11.996 |  | 2.170 |  | 1.310 |  | 3.030 |  | < .001 | *** |
|  |  | sync, hard, group correct |  | 0.218 |  | 0.146 |  | 0.290 |  | 0.023 |  | 9.557 |  | 1.680 |  | 0.935 |  | 2.426 |  | < .001 | *** |
| sync, easy, group incorrect |  | async, hard, group incorrect |  | 0.385 |  | 0.316 |  | 0.454 |  | 0.022 |  | 17.699 |  | 2.967 |  | 1.939 |  | 3.994 |  | < .001 | *** |
|  |  | sync, hard, group incorrect |  | 0.398 |  | 0.330 |  | 0.466 |  | 0.022 |  | 18.503 |  | 3.070 |  | 2.018 |  | 4.121 |  | < .001 | *** |
|  |  | async, easy, group correct |  | -0.014 |  | -0.090 |  | 0.063 |  | 0.024 |  | -0.557 |  | -0.104 |  | -0.693 |  | 0.485 |  | 1.000 |  |
|  |  | sync, easy, group correct |  | -0.010 |  | -0.087 |  | 0.067 |  | 0.024 |  | -0.411 |  | -0.077 |  | -0.670 |  | 0.516 |  | 1.000 |  |
|  |  | async, hard, group correct |  | 0.305 |  | 0.232 |  | 0.377 |  | 0.023 |  | 13.362 |  | 2.349 |  | 1.459 |  | 3.240 |  | < .001 | *** |
|  |  | sync, hard, group correct |  | 0.241 |  | 0.167 |  | 0.315 |  | 0.023 |  | 10.281 |  | 1.860 |  | 1.066 |  | 2.653 |  | < .001 | *** |
| async, hard, group incorrect |  | sync, hard, group incorrect |  | 0.013 |  | -0.051 |  | 0.078 |  | 0.020 |  | 0.655 |  | 0.103 |  | -0.392 |  | 0.598 |  | 1.000 |  |
|  |  | async, easy, group correct |  | -0.398 |  | -0.473 |  | -0.324 |  | 0.023 |  | -16.976 |  | -3.071 |  | -4.146 |  | -1.995 |  | < .001 | *** |
|  |  | sync, easy, group correct |  | -0.395 |  | -0.467 |  | -0.323 |  | 0.023 |  | -17.312 |  | -3.044 |  | -4.105 |  | -1.984 |  | < .001 | *** |
|  |  | async, hard, group correct |  | -0.080 |  | -0.157 |  | -0.003 |  | 0.024 |  | -3.278 |  | -0.617 |  | -1.238 |  | 0.003 |  | 0.011 | * |
|  |  | sync, hard, group correct |  | -0.144 |  | -0.220 |  | -0.067 |  | 0.024 |  | -5.921 |  | -1.107 |  | -1.781 |  | -0.433 |  | < .001 | *** |
| sync, hard, group incorrect |  | async, easy, group correct |  | -0.412 |  | -0.484 |  | -0.339 |  | 0.023 |  | -18.049 |  | -3.174 |  | -4.267 |  | -2.080 |  | < .001 | *** |
|  |  | sync, easy, group correct |  | -0.408 |  | -0.482 |  | -0.334 |  | 0.023 |  | -17.396 |  | -3.147 |  | -4.242 |  | -2.052 |  | < .001 | *** |
|  |  | async, hard, group correct |  | -0.093 |  | -0.170 |  | -0.017 |  | 0.024 |  | -3.852 |  | -0.720 |  | -1.346 |  | -0.094 |  | 0.002 | ** |
|  |  | sync, hard, group correct |  | -0.157 |  | -0.234 |  | -0.080 |  | 0.024 |  | -6.424 |  | -1.210 |  | -1.903 |  | -0.517 |  | < .001 | *** |
| async, easy, group correct |  | sync, easy, group correct |  | 0.003 |  | -0.061 |  | 0.068 |  | 0.020 |  | 0.170 |  | 0.027 |  | -0.467 |  | 0.521 |  | 1.000 |  |
|  |  | async, hard, group correct |  | 0.318 |  | 0.250 |  | 0.386 |  | 0.022 |  | 14.789 |  | 2.454 |  | 1.557 |  | 3.350 |  | < .001 | *** |
|  |  | sync, hard, group correct |  | 0.255 |  | 0.186 |  | 0.323 |  | 0.022 |  | 11.717 |  | 1.964 |  | 1.177 |  | 2.751 |  | < .001 | *** |
| sync, easy, group correct |  | async, hard, group correct |  | 0.315 |  | 0.246 |  | 0.383 |  | 0.022 |  | 14.478 |  | 2.427 |  | 1.533 |  | 3.320 |  | < .001 | *** |
|  |  | sync, hard, group correct |  | 0.251 |  | 0.183 |  | 0.319 |  | 0.022 |  | 11.676 |  | 1.937 |  | 1.160 |  | 2.714 |  | < .001 | *** |
| async, hard, group correct |  | sync, hard, group correct |  | -0.064 |  | -0.128 |  | 8.610×10^-4^ |  | 0.020 |  | -3.120 |  | -0.490 |  | -1.004 |  | 0.025 |  | 0.016 | * |
| * p < .05, ** p < .01, *** p < .001 | | | | | | | | | | | | | | | | | | | | | |
| *Note.*  P-value and confidence intervals adjusted for comparing a family of 28 estimates (confidence intervals corrected using the bonferroni method). | | | | | | | | | | | | | | | | | | | | | |
| *Note.*  Results are averaged over the levels of: Blockfirst  **Table S7.** Summary of the post-hoc pairwise comparisons for accuracy across block order (Blockfirst) and synchrony conditions. The table presents mean differences, 95% CI, SE, t-values, Cohen’s d, and multiple comparison adjustments. | | | | | | | | | | | | | | | | | | | | | |

| *Post Hoc Comparisons - Blockfirst ✻ Synchrony* | | | | | | | | | | | | | | | | | | | | | |
| --- | --- | --- | --- | --- | --- | --- | --- | --- | --- | --- | --- | --- | --- | --- | --- | --- | --- | --- | --- | --- | --- |
|  | | | | | | 95% CI for Mean Difference | | | |  | | | | | | 95% CI for Cohen's d | | | |  | |
|  | |  | | Mean Difference | | Lower | | Upper | | SE | | t | | Cohen's d | | Lower | | Upper | | p_holm_ | |
| AsyncFirst, async |  | SyncFirst, async |  | -0.040 |  | -0.091 |  | 0.011 |  | 0.019 |  | -2.112 |  | -0.307 |  | -0.706 |  | 0.093 |  | 0.187 |  |
|  |  | AsyncFirst, sync |  | -0.040 |  | -0.078 |  | -0.002 |  | 0.014 |  | -2.879 |  | -0.308 |  | -0.607 |  | -0.009 |  | 0.034 | * |
|  |  | SyncFirst, sync |  | -0.035 |  | -0.086 |  | 0.016 |  | 0.019 |  | -1.848 |  | -0.269 |  | -0.666 |  | 0.129 |  | 0.271 |  |
| SyncFirst, async |  | AsyncFirst, sync |  | -1.541×10^-4^ |  | -0.051 |  | 0.051 |  | 0.019 |  | -0.008 |  | -0.001 |  | -0.393 |  | 0.391 |  | 1.000 |  |
|  |  | SyncFirst, sync |  | 0.005 |  | -0.033 |  | 0.043 |  | 0.014 |  | 0.358 |  | 0.038 |  | -0.250 |  | 0.327 |  | 1.000 |  |
| AsyncFirst, sync |  | SyncFirst, sync |  | 0.005 |  | -0.046 |  | 0.056 |  | 0.019 |  | 0.272 |  | 0.039 |  | -0.352 |  | 0.431 |  | 1.000 |  |
| * p < .05, *** p < .001 | | | | | | | | | | | | | | | | | | | | | |
| *Note.*  P-value and confidence intervals adjusted for comparing a family of 6 estimates (confidence intervals corrected using the bonferroni method). | | | | | | | | | | | | | | | | | | | | | |
| *Note.*  Results are averaged over the levels of: Difficulty, Group Correctness d) **Inverse Efficiency Scores– PostHoc Pairwairse Comparisons** **Table S8.** Summary of the post-hoc pairwise comparisons for inverse efficiency scores across difficulty and group correctness conditions. The table presents mean differences, 95% CI, SE, t-values, Cohen’s d, and multiple comparison adjustments. *Post Hoc Comparisons - Difficulty ✻ Group Correctness* \| \| \| \| \| \| \| \| \| \| \| \| \| \| \| \| \| \| \| \| \|; \| \| \| \| \| \| 95% CI for Mean Difference \| \| \| \| \| \| \| \| \| \| 95% CI for Cohen's d \| \| \| \| \|; \| \| \| \| Mean Difference \| \| Lower \| \| Upper \| \| SE \| \| t \| \| Cohen's d \| \| Lower \| \| Upper \| \| p_holm_ \|; easy, group incorrect \| \| hard, group incorrect \| \| -0.853 \| \| -1.045 \| \| -0.661 \| \| 0.071 \| \| -11.941 \| \| -1.516 \| \| -2.036 \| \| -0.995 \| \| < .001 \| ***; \| \| easy, group correct \| \| 0.034 \| \| -0.138 \| \| 0.205 \| \| 0.064 \| \| 0.526 \| \| 0.060 \| \| -0.244 \| \| 0.364 \| \| 0.600 \|; \| \| hard, group correct \| \| -0.482 \| \| -0.682 \| \| -0.282 \| \| 0.074 \| \| -6.488 \| \| -0.857 \| \| -1.275 \| \| -0.439 \| \| < .001 \| ***; hard, group incorrect \| \| easy, group correct \| \| 0.886 \| \| 0.687 \| \| 1.086 \| \| 0.074 \| \| 11.931 \| \| 1.575 \| \| 1.034 \| \| 2.116 \| \| < .001 \| ***; \| \| hard, group correct \| \| 0.371 \| \| 0.199 \| \| 0.542 \| \| 0.064 \| \| 5.812 \| \| 0.659 \| \| 0.310 \| \| 1.008 \| \| < .001 \| ***; easy, group correct \| \| hard, group correct \| \| -0.516 \| \| -0.708 \| \| -0.323 \| \| 0.071 \| \| -7.219 \| \| -0.916 \| \| -1.331 \| \| -0.501 \| \| < .001 \| ***; *** p < .001 \| \| \| \| \| \| \| \| \| \| \| \| \| \| \| \| \| \| \| \| \|; Note.  P-value and confidence intervals adjusted for comparing a family of 6 estimates (confidence intervals corrected using the bonferroni method). \| \| \| \| \| \| \| \| \| \| \| \| \| \| \| \| \| \| \| \| \|; Note.  Results are averaged over the levels of: Blockfirst, Synchrony \| \| \| \| \| \| \| \| \| \| \| \| \| \| \| \| \| \| \| \| \| | | | | | | | | | | | | | | | | | | | | | |

**Table S9.** Summary of the post-hoc pairwise comparisons for inverse efficiency scores across synchrony and group correctness conditions. The table provides statistical values including mean differences, 95% CI, SE, t-values, Cohen’s d, and corrected p-values.

| *Post Hoc Comparisons - Synchrony ✻ Group Correctness* | | | | | | | | | | | | | | | | | | | | | |
| --- | --- | --- | --- | --- | --- | --- | --- | --- | --- | --- | --- | --- | --- | --- | --- | --- | --- | --- | --- | --- | --- |
|  | | | | | | 95% CI for Mean Difference | | | |  | | | | | | 95% CI for Cohen's d | | | |  | |
|  | |  | | Mean Difference | | Lower | | Upper | | SE | | t | | Cohen's d | | Lower | | Upper | | p_holm_ | |
| async, group incorrect |  | sync, group incorrect |  | 0.027 |  | -0.129 |  | 0.182 |  | 0.058 |  | 0.463 |  | 0.047 |  | -0.228 |  | 0.323 |  | 1.000 |  |
|  |  | async, group correct |  | 0.198 |  | 0.047 |  | 0.349 |  | 0.056 |  | 3.529 |  | 0.352 |  | 0.069 |  | 0.634 |  | 0.003 | ** |
|  |  | sync, group correct |  | 0.233 |  | 0.049 |  | 0.417 |  | 0.068 |  | 3.413 |  | 0.414 |  | 0.071 |  | 0.758 |  | 0.004 | ** |
| sync, group incorrect |  | async, group correct |  | 0.171 |  | -0.013 |  | 0.355 |  | 0.068 |  | 2.503 |  | 0.304 |  | -0.032 |  | 0.640 |  | 0.041 | * |
|  |  | sync, group correct |  | 0.207 |  | 0.055 |  | 0.358 |  | 0.056 |  | 3.684 |  | 0.367 |  | 0.083 |  | 0.651 |  | 0.002 | ** |
| async, group correct |  | sync, group correct |  | 0.035 |  | -0.120 |  | 0.191 |  | 0.058 |  | 0.614 |  | 0.063 |  | -0.213 |  | 0.339 |  | 1.000 |  |
| * p < .05, ** p < .01, *** p < .001 | | | | | | | | | | | | | | | | | | | | | |
| *Note.*  P-value and confidence intervals adjusted for comparing a family of 6 estimates (confidence intervals corrected using the bonferroni method). | | | | | | | | | | | | | | | | | | | | | |
| *Note.*  Results are averaged over the levels of: Blockfirst, Difficulty  **Table S10.** Summary of the post-hoc pairwise comparisons for inverse efficiency scores across synchrony, difficulty, and group correctness conditions. The table includes statistical comparisons such as mean differences, 95% CI, SE, t-values, Cohen’s d, and adjusted p-values. | | | | | | | | | | | | | | | | | | | | | |

| *Post Hoc Comparisons - Synchrony ✻ Difficulty ✻ Group Correctness* | | | | | | | | | | | | | | | | | | | | | |
| --- | --- | --- | --- | --- | --- | --- | --- | --- | --- | --- | --- | --- | --- | --- | --- | --- | --- | --- | --- | --- | --- |
|  | | | | | | 95% CI for Mean Difference | | | |  | | | | | | 95% CI for Cohen's d | | | |  | |
|  | |  | | Mean Difference | | Lower | | Upper | | SE | | t | | Cohen's d | | Lower | | Upper | | p_holm_ | |
| async, easy, group incorrect |  | sync, easy, group incorrect |  | 0.057 |  | -0.181 |  | 0.296 |  | 0.075 |  | 0.763 |  | 0.102 |  | -0.321 |  | 0.525 |  | 1.000 |  |
|  |  | async, hard, group incorrect |  | -0.822 |  | -1.096 |  | -0.549 |  | 0.086 |  | -9.550 |  | -1.461 |  | -2.121 |  | -0.801 |  | < .001 | *** |
|  |  | sync, hard, group incorrect |  | -0.826 |  | -1.117 |  | -0.535 |  | 0.092 |  | -8.999 |  | -1.468 |  | -2.153 |  | -0.783 |  | < .001 | *** |
|  |  | async, easy, group correct |  | 0.060 |  | -0.180 |  | 0.301 |  | 0.076 |  | 0.792 |  | 0.107 |  | -0.321 |  | 0.534 |  | 1.000 |  |
|  |  | sync, easy, group correct |  | 0.064 |  | -0.219 |  | 0.348 |  | 0.090 |  | 0.719 |  | 0.114 |  | -0.390 |  | 0.619 |  | 1.000 |  |
|  |  | async, hard, group correct |  | -0.487 |  | -0.769 |  | -0.204 |  | 0.089 |  | -5.460 |  | -0.865 |  | -1.432 |  | -0.298 |  | < .001 | *** |
|  |  | sync, hard, group correct |  | -0.420 |  | -0.716 |  | -0.124 |  | 0.093 |  | -4.496 |  | -0.747 |  | -1.320 |  | -0.173 |  | < .001 | *** |
| sync, easy, group incorrect |  | async, hard, group incorrect |  | -0.880 |  | -1.171 |  | -0.589 |  | 0.092 |  | -9.582 |  | -1.563 |  | -2.268 |  | -0.858 |  | < .001 | *** |
|  |  | sync, hard, group incorrect |  | -0.883 |  | -1.157 |  | -0.610 |  | 0.086 |  | -10.260 |  | -1.570 |  | -2.253 |  | -0.887 |  | < .001 | *** |
|  |  | async, easy, group correct |  | 0.003 |  | -0.281 |  | 0.286 |  | 0.090 |  | 0.031 |  | 0.005 |  | -0.499 |  | 0.508 |  | 1.000 |  |
|  |  | sync, easy, group correct |  | 0.007 |  | -0.233 |  | 0.248 |  | 0.076 |  | 0.093 |  | 0.013 |  | -0.414 |  | 0.439 |  | 1.000 |  |
|  |  | async, hard, group correct |  | -0.544 |  | -0.840 |  | -0.248 |  | 0.093 |  | -5.821 |  | -0.967 |  | -1.570 |  | -0.363 |  | < .001 | *** |
|  |  | sync, hard, group correct |  | -0.477 |  | -0.760 |  | -0.195 |  | 0.089 |  | -5.356 |  | -0.848 |  | -1.413 |  | -0.284 |  | < .001 | *** |
| async, hard, group incorrect |  | sync, hard, group incorrect |  | -0.004 |  | -0.242 |  | 0.234 |  | 0.075 |  | -0.051 |  | -0.007 |  | -0.429 |  | 0.415 |  | 1.000 |  |
|  |  | async, easy, group correct |  | 0.882 |  | 0.600 |  | 1.165 |  | 0.089 |  | 9.899 |  | 1.568 |  | 0.873 |  | 2.263 |  | < .001 | *** |
|  |  | sync, easy, group correct |  | 0.887 |  | 0.590 |  | 1.183 |  | 0.093 |  | 9.488 |  | 1.576 |  | 0.861 |  | 2.290 |  | < .001 | *** |
|  |  | async, hard, group correct |  | 0.336 |  | 0.095 |  | 0.576 |  | 0.076 |  | 4.425 |  | 0.596 |  | 0.132 |  | 1.060 |  | < .001 | *** |
|  |  | sync, hard, group correct |  | 0.402 |  | 0.119 |  | 0.686 |  | 0.090 |  | 4.489 |  | 0.715 |  | 0.165 |  | 1.264 |  | < .001 | *** |
| sync, hard, group incorrect |  | async, easy, group correct |  | 0.886 |  | 0.590 |  | 1.182 |  | 0.093 |  | 9.483 |  | 1.575 |  | 0.861 |  | 2.289 |  | < .001 | *** |
|  |  | sync, easy, group correct |  | 0.890 |  | 0.608 |  | 1.173 |  | 0.089 |  | 9.990 |  | 1.582 |  | 0.884 |  | 2.280 |  | < .001 | *** |
|  |  | async, hard, group correct |  | 0.339 |  | 0.056 |  | 0.623 |  | 0.090 |  | 3.790 |  | 0.603 |  | 0.067 |  | 1.140 |  | 0.002 | ** |
|  |  | sync, hard, group correct |  | 0.406 |  | 0.165 |  | 0.647 |  | 0.076 |  | 5.353 |  | 0.721 |  | 0.241 |  | 1.202 |  | < .001 | *** |
| async, easy, group correct |  | sync, easy, group correct |  | 0.004 |  | -0.234 |  | 0.243 |  | 0.075 |  | 0.057 |  | 0.008 |  | -0.414 |  | 0.430 |  | 1.000 |  |
|  |  | async, hard, group correct |  | -0.547 |  | -0.820 |  | -0.273 |  | 0.086 |  | -6.350 |  | -0.972 |  | -1.540 |  | -0.403 |  | < .001 | *** |
|  |  | sync, hard, group correct |  | -0.480 |  | -0.771 |  | -0.189 |  | 0.092 |  | -5.231 |  | -0.853 |  | -1.432 |  | -0.275 |  | < .001 | *** |
| sync, easy, group correct |  | async, hard, group correct |  | -0.551 |  | -0.842 |  | -0.260 |  | 0.092 |  | -6.002 |  | -0.979 |  | -1.576 |  | -0.382 |  | < .001 | *** |
|  |  | sync, hard, group correct |  | -0.484 |  | -0.758 |  | -0.211 |  | 0.086 |  | -5.627 |  | -0.861 |  | -1.412 |  | -0.309 |  | < .001 | *** |
| async, hard, group correct |  | sync, hard, group correct |  | 0.067 |  | -0.172 |  | 0.305 |  | 0.075 |  | 0.886 |  | 0.118 |  | -0.305 |  | 0.542 |  | 1.000 |  |
| ** p < .01, *** p < .001 | | | | | | | | | | | | | | | | | | | | | |
| *Note.*  P-value and confidence intervals adjusted for comparing a family of 28 estimates (confidence intervals corrected using the bonferroni method). | | | | | | | | | | | | | | | | | | | | | |
| *Note.*  Results are averaged over the levels of: Blockfirst | | | | | | | | | | | | | | | | | | | | | |

**Table S11.** Summary of the post-hoc pairwise comparisons for inverse efficiency scores across synchrony and block order (Blockfirst). The table includes statistical comparisons such as mean differences, 95% CI, SE, t-values, Cohen’s d, and adjusted p-values.

| *Post Hoc Comparisons - Blockfirst ✻ Synchrony* | | | | | | | | | | | | | | | | | | | | | |
| --- | --- | --- | --- | --- | --- | --- | --- | --- | --- | --- | --- | --- | --- | --- | --- | --- | --- | --- | --- | --- | --- |
|  | | | | | | 95% CI for Mean Difference | | | |  | | | | | | 95% CI for Cohen's d | | | |  | |
|  | |  | | Mean Difference | | Lower | | Upper | | SE | | t | | Cohen's d | | Lower | | Upper | | p_holm_ | |
| AsyncFirst, async |  | SyncFirst, async |  | 0.226 |  | -0.077 |  | 0.529 |  | 0.112 |  | 2.018 |  | 0.402 |  | -0.148 |  | 0.951 |  | 0.188 |  |
|  |  | AsyncFirst, sync |  | 0.250 |  | 0.054 |  | 0.446 |  | 0.072 |  | 3.494 |  | 0.444 |  | 0.081 |  | 0.808 |  | 0.006 | ** |
|  |  | SyncFirst, sync |  | 0.038 |  | -0.265 |  | 0.341 |  | 0.112 |  | 0.340 |  | 0.068 |  | -0.472 |  | 0.607 |  | 1.000 |  |
| SyncFirst, async |  | AsyncFirst, sync |  | 0.024 |  | -0.279 |  | 0.327 |  | 0.112 |  | 0.214 |  | 0.043 |  | -0.497 |  | 0.582 |  | 1.000 |  |
|  |  | SyncFirst, sync |  | -0.188 |  | -0.374 |  | -0.002 |  | 0.068 |  | -2.773 |  | -0.334 |  | -0.672 |  | 0.004 |  | 0.038 | * |
| AsyncFirst, sync |  | SyncFirst, sync |  | -0.212 |  | -0.515 |  | 0.091 |  | 0.112 |  | -1.892 |  | -0.377 |  | -0.925 |  | 0.172 |  | 0.188 |  |
| * p < .05, ** p < .01, *** p < .001 | | | | | | | | | | | | | | | | | | | | | |
| *Note.*  P-value and confidence intervals adjusted for comparing a family of 6 estimates (confidence intervals corrected using the bonferroni method). | | | | | | | | | | | | | | | | | | | | | |
| *Note.*  Results are averaged over the levels of: Difficulty, Group Correctness  **Table S12.** Summary of the post-hoc pairwise comparisons for block order (Blockfirst), synchrony, difficulty, and group correctness. The table reports the mean difference, 95% CI, SE, t-values, Cohen’s d, and multiple comparison p-value adjustments. | | | | | | | | | | | | | | | | | | | | | |

| *Post Hoc Comparisons - Blockfirst ✻ Synchrony ✻ Group Correctness* | | | | | | | | | | | | | | | | | | | | | |
| --- | --- | --- | --- | --- | --- | --- | --- | --- | --- | --- | --- | --- | --- | --- | --- | --- | --- | --- | --- | --- | --- |
|  | | | | | | 95% CI for Mean Difference | | | |  | | | | | | 95% CI for Cohen's d | | | |  | |
|  | |  | | Mean Difference | | Lower | | Upper | | SE | | t | | Cohen's d | | Lower | | Upper | | p_holm_ | |
| AsyncFirst, async, group incorrect |  | SyncFirst, async, group incorrect |  | 0.264 |  | -0.137 |  | 0.664 |  | 0.125 |  | 2.104 |  | 0.468 |  | -0.259 |  | 1.195 |  | 0.638 |  |
|  |  | AsyncFirst, sync, group incorrect |  | 0.320 |  | 0.050 |  | 0.590 |  | 0.084 |  | 3.824 |  | 0.569 |  | 0.062 |  | 1.077 |  | 0.006 | ** |
|  |  | SyncFirst, sync, group incorrect |  | -0.003 |  | -0.404 |  | 0.397 |  | 0.125 |  | -0.026 |  | -0.006 |  | -0.718 |  | 0.707 |  | 1.000 |  |
|  |  | AsyncFirst, async, group correct |  | 0.235 |  | -0.027 |  | 0.498 |  | 0.081 |  | 2.891 |  | 0.418 |  | -0.062 |  | 0.899 |  | 0.101 |  |
|  |  | SyncFirst, async, group correct |  | 0.424 |  | 0.023 |  | 0.825 |  | 0.125 |  | 3.384 |  | 0.753 |  | 0.003 |  | 1.503 |  | 0.023 | * |
|  |  | AsyncFirst, sync, group correct |  | 0.415 |  | 0.097 |  | 0.733 |  | 0.099 |  | 4.182 |  | 0.738 |  | 0.129 |  | 1.346 |  | 0.002 | ** |
|  |  | SyncFirst, sync, group correct |  | 0.315 |  | -0.086 |  | 0.716 |  | 0.125 |  | 2.514 |  | 0.560 |  | -0.174 |  | 1.293 |  | 0.253 |  |
| SyncFirst, async, group incorrect |  | AsyncFirst, sync, group incorrect |  | 0.057 |  | -0.344 |  | 0.457 |  | 0.125 |  | 0.453 |  | 0.101 |  | -0.612 |  | 0.814 |  | 1.000 |  |
|  |  | SyncFirst, sync, group incorrect |  | -0.267 |  | -0.522 |  | -0.011 |  | 0.079 |  | -3.365 |  | -0.474 |  | -0.949 |  | 3.760×10^-4^ |  | 0.026 | * |
|  |  | AsyncFirst, async, group correct |  | -0.028 |  | -0.429 |  | 0.372 |  | 0.125 |  | -0.225 |  | -0.050 |  | -0.763 |  | 0.662 |  | 1.000 |  |
|  |  | SyncFirst, async, group correct |  | 0.160 |  | -0.088 |  | 0.409 |  | 0.077 |  | 2.080 |  | 0.285 |  | -0.162 |  | 0.732 |  | 0.647 |  |
|  |  | AsyncFirst, sync, group correct |  | 0.152 |  | -0.249 |  | 0.552 |  | 0.125 |  | 1.210 |  | 0.269 |  | -0.448 |  | 0.987 |  | 1.000 |  |
|  |  | SyncFirst, sync, group correct |  | 0.051 |  | -0.250 |  | 0.353 |  | 0.094 |  | 0.546 |  | 0.091 |  | -0.444 |  | 0.627 |  | 1.000 |  |
| AsyncFirst, sync, group incorrect |  | SyncFirst, sync, group incorrect |  | -0.324 |  | -0.724 |  | 0.077 |  | 0.125 |  | -2.583 |  | -0.575 |  | -1.309 |  | 0.160 |  | 0.221 |  |
|  |  | AsyncFirst, async, group correct |  | -0.085 |  | -0.403 |  | 0.233 |  | 0.099 |  | -0.856 |  | -0.151 |  | -0.717 |  | 0.415 |  | 1.000 |  |
|  |  | SyncFirst, async, group correct |  | 0.104 |  | -0.297 |  | 0.504 |  | 0.125 |  | 0.827 |  | 0.184 |  | -0.531 |  | 0.899 |  | 1.000 |  |
|  |  | AsyncFirst, sync, group correct |  | 0.095 |  | -0.167 |  | 0.357 |  | 0.081 |  | 1.165 |  | 0.169 |  | -0.297 |  | 0.634 |  | 1.000 |  |
|  |  | SyncFirst, sync, group correct |  | -0.005 |  | -0.406 |  | 0.395 |  | 0.125 |  | -0.043 |  | -0.010 |  | -0.722 |  | 0.703 |  | 1.000 |  |
| SyncFirst, sync, group incorrect |  | AsyncFirst, async, group correct |  | 0.239 |  | -0.162 |  | 0.639 |  | 0.125 |  | 1.905 |  | 0.424 |  | -0.300 |  | 1.149 |  | 0.889 |  |
|  |  | SyncFirst, async, group correct |  | 0.427 |  | 0.126 |  | 0.728 |  | 0.094 |  | 4.545 |  | 0.759 |  | 0.175 |  | 1.343 |  | < .001 | *** |
|  |  | AsyncFirst, sync, group correct |  | 0.418 |  | 0.018 |  | 0.819 |  | 0.125 |  | 3.341 |  | 0.744 |  | -0.005 |  | 1.493 |  | 0.026 | * |
|  |  | SyncFirst, sync, group correct |  | 0.318 |  | 0.070 |  | 0.566 |  | 0.077 |  | 4.128 |  | 0.565 |  | 0.093 |  | 1.038 |  | 0.002 | ** |
| AsyncFirst, async, group correct |  | SyncFirst, async, group correct |  | 0.188 |  | -0.212 |  | 0.589 |  | 0.125 |  | 1.505 |  | 0.335 |  | -0.385 |  | 1.055 |  | 1.000 |  |
|  |  | AsyncFirst, sync, group correct |  | 0.180 |  | -0.090 |  | 0.450 |  | 0.084 |  | 2.147 |  | 0.319 |  | -0.167 |  | 0.805 |  | 0.623 |  |
|  |  | SyncFirst, sync, group correct |  | 0.080 |  | -0.321 |  | 0.480 |  | 0.125 |  | 0.635 |  | 0.141 |  | -0.572 |  | 0.855 |  | 1.000 |  |
| SyncFirst, async, group correct |  | AsyncFirst, sync, group correct |  | -0.009 |  | -0.409 |  | 0.392 |  | 0.125 |  | -0.070 |  | -0.015 |  | -0.728 |  | 0.697 |  | 1.000 |  |
|  |  | SyncFirst, sync, group correct |  | -0.109 |  | -0.364 |  | 0.147 |  | 0.079 |  | -1.374 |  | -0.194 |  | -0.649 |  | 0.261 |  | 1.000 |  |
| AsyncFirst, sync, group correct |  | SyncFirst, sync, group correct |  | -0.100 |  | -0.501 |  | 0.300 |  | 0.125 |  | -0.800 |  | -0.178 |  | -0.893 |  | 0.536 |  | 1.000 |  |
| * p < .05, ** p < .01, *** p < .001 | | | | | | | | | | | | | | | | | | | | | |
| *Note.*  P-value and confidence intervals adjusted for comparing a family of 28 estimates (confidence intervals corrected using the bonferroni method). | | | | | | | | | | | | | | | | | | | | | |
| *Note.*  Results are averaged over the levels of: Difficulty | | | | | | | | | | | | | | | | | | | | | |

| *Post Hoc Comparisons - Blockfirst ✻ Synchrony ✻ Difficulty ✻ Group Correctness* | | | | | | | | | | | | | | | | | | | | | |
| --- | --- | --- | --- | --- | --- | --- | --- | --- | --- | --- | --- | --- | --- | --- | --- | --- | --- | --- | --- | --- | --- |
|  | | | | | | 95% CI for Mean Difference | | | |  | | | | | | 95% CI for Cohen's d | | | |  | |
|  | |  | | Mean Difference | | Lower | | Upper | | SE | | t | | Cohen's d | | Lower | | Upper | | p_holm_ | |
| AsyncFirst, async, easy, group incorrect |  | SyncFirst, async, easy, group incorrect |  | 0.106 |  | -0.439 |  | 0.651 |  | 0.152 |  | 0.697 |  | 0.188 |  | -0.783 |  | 1.159 |  | 1.000 |  |
|  |  | AsyncFirst, sync, easy, group incorrect |  | 0.184 |  | -0.209 |  | 0.576 |  | 0.109 |  | 1.685 |  | 0.327 |  | -0.377 |  | 1.030 |  | 1.000 |  |
|  |  | SyncFirst, sync, easy, group incorrect |  | 0.037 |  | -0.508 |  | 0.582 |  | 0.152 |  | 0.242 |  | 0.065 |  | -0.904 |  | 1.035 |  | 1.000 |  |
|  |  | AsyncFirst, async, hard, group incorrect |  | -0.980 |  | -1.430 |  | -0.530 |  | 0.125 |  | -7.836 |  | -1.741 |  | -2.742 |  | -0.741 |  | < .001 | *** |
|  |  | SyncFirst, async, hard, group incorrect |  | -0.559 |  | -1.104 |  | -0.014 |  | 0.152 |  | -3.676 |  | -0.993 |  | -2.022 |  | 0.036 |  | 0.020 | * |
|  |  | AsyncFirst, sync, hard, group incorrect |  | -0.523 |  | -1.002 |  | -0.044 |  | 0.133 |  | -3.923 |  | -0.929 |  | -1.838 |  | -0.021 |  | 0.009 | ** |
|  |  | SyncFirst, sync, hard, group incorrect |  | -1.023 |  | -1.568 |  | -0.478 |  | 0.152 |  | -6.733 |  | -1.818 |  | -2.976 |  | -0.661 |  | < .001 | *** |
|  |  | AsyncFirst, async, easy, group correct |  | 0.045 |  | -0.351 |  | 0.441 |  | 0.110 |  | 0.410 |  | 0.080 |  | -0.622 |  | 0.782 |  | 1.000 |  |
|  |  | SyncFirst, async, easy, group correct |  | 0.181 |  | -0.364 |  | 0.726 |  | 0.152 |  | 1.190 |  | 0.321 |  | -0.654 |  | 1.297 |  | 1.000 |  |
|  |  | AsyncFirst, sync, easy, group correct |  | 0.214 |  | -0.253 |  | 0.680 |  | 0.130 |  | 1.643 |  | 0.380 |  | -0.459 |  | 1.219 |  | 1.000 |  |
|  |  | SyncFirst, sync, easy, group correct |  | 0.021 |  | -0.524 |  | 0.566 |  | 0.152 |  | 0.138 |  | 0.037 |  | -0.932 |  | 1.006 |  | 1.000 |  |
|  |  | AsyncFirst, async, hard, group correct |  | -0.554 |  | -1.020 |  | -0.089 |  | 0.129 |  | -4.282 |  | -0.985 |  | -1.878 |  | -0.092 |  | 0.002 | ** |
|  |  | SyncFirst, async, hard, group correct |  | -0.313 |  | -0.858 |  | 0.232 |  | 0.152 |  | -2.060 |  | -0.556 |  | -1.544 |  | 0.432 |  | 1.000 |  |
|  |  | AsyncFirst, sync, hard, group correct |  | -0.363 |  | -0.851 |  | 0.125 |  | 0.136 |  | -2.677 |  | -0.646 |  | -1.538 |  | 0.247 |  | 0.413 |  |
|  |  | SyncFirst, sync, hard, group correct |  | -0.371 |  | -0.916 |  | 0.174 |  | 0.152 |  | -2.441 |  | -0.659 |  | -1.655 |  | 0.336 |  | 0.726 |  |
| SyncFirst, async, easy, group incorrect |  | AsyncFirst, sync, easy, group incorrect |  | 0.078 |  | -0.467 |  | 0.623 |  | 0.152 |  | 0.512 |  | 0.138 |  | -0.832 |  | 1.108 |  | 1.000 |  |
|  |  | SyncFirst, sync, easy, group incorrect |  | -0.069 |  | -0.441 |  | 0.302 |  | 0.103 |  | -0.670 |  | -0.123 |  | -0.782 |  | 0.537 |  | 1.000 |  |
|  |  | AsyncFirst, async, hard, group incorrect |  | -1.086 |  | -1.631 |  | -0.541 |  | 0.152 |  | -7.144 |  | -1.930 |  | -3.109 |  | -0.750 |  | < .001 | *** |
|  |  | SyncFirst, async, hard, group incorrect |  | -0.665 |  | -1.091 |  | -0.238 |  | 0.118 |  | -5.614 |  | -1.181 |  | -2.041 |  | -0.321 |  | < .001 | *** |
|  |  | AsyncFirst, sync, hard, group incorrect |  | -0.629 |  | -1.174 |  | -0.084 |  | 0.152 |  | -4.138 |  | -1.118 |  | -2.162 |  | -0.074 |  | 0.004 | ** |
|  |  | SyncFirst, sync, hard, group incorrect |  | -1.129 |  | -1.583 |  | -0.675 |  | 0.126 |  | -8.945 |  | -2.007 |  | -3.073 |  | -0.941 |  | < .001 | *** |
|  |  | AsyncFirst, async, easy, group correct |  | -0.061 |  | -0.606 |  | 0.484 |  | 0.152 |  | -0.400 |  | -0.108 |  | -1.078 |  | 0.862 |  | 1.000 |  |
|  |  | SyncFirst, async, easy, group correct |  | 0.075 |  | -0.300 |  | 0.450 |  | 0.104 |  | 0.718 |  | 0.133 |  | -0.533 |  | 0.800 |  | 1.000 |  |
|  |  | AsyncFirst, sync, easy, group correct |  | 0.108 |  | -0.437 |  | 0.653 |  | 0.152 |  | 0.709 |  | 0.192 |  | -0.780 |  | 1.163 |  | 1.000 |  |
|  |  | SyncFirst, sync, easy, group correct |  | -0.085 |  | -0.527 |  | 0.357 |  | 0.123 |  | -0.690 |  | -0.151 |  | -0.938 |  | 0.636 |  | 1.000 |  |
|  |  | AsyncFirst, async, hard, group correct |  | -0.660 |  | -1.205 |  | -0.115 |  | 0.152 |  | -4.344 |  | -1.173 |  | -2.225 |  | -0.122 |  | 0.002 | ** |
|  |  | SyncFirst, async, hard, group correct |  | -0.419 |  | -0.860 |  | 0.022 |  | 0.123 |  | -3.418 |  | -0.745 |  | -1.568 |  | 0.079 |  | 0.048 | * |
|  |  | AsyncFirst, sync, hard, group correct |  | -0.469 |  | -1.014 |  | 0.076 |  | 0.152 |  | -3.088 |  | -0.834 |  | -1.845 |  | 0.178 |  | 0.130 |  |
|  |  | SyncFirst, sync, hard, group correct |  | -0.477 |  | -0.939 |  | -0.015 |  | 0.128 |  | -3.712 |  | -0.848 |  | -1.718 |  | 0.023 |  | 0.019 | * |
| AsyncFirst, sync, easy, group incorrect |  | SyncFirst, sync, easy, group incorrect |  | -0.147 |  | -0.692 |  | 0.398 |  | 0.152 |  | -0.967 |  | -0.261 |  | -1.234 |  | 0.712 |  | 1.000 |  |
|  |  | AsyncFirst, async, hard, group incorrect |  | -1.164 |  | -1.643 |  | -0.684 |  | 0.133 |  | -8.728 |  | -2.068 |  | -3.181 |  | -0.955 |  | < .001 | *** |
|  |  | SyncFirst, async, hard, group incorrect |  | -0.742 |  | -1.287 |  | -0.198 |  | 0.152 |  | -4.885 |  | -1.319 |  | -2.392 |  | -0.247 |  | < .001 | *** |
|  |  | AsyncFirst, sync, hard, group incorrect |  | -0.707 |  | -1.157 |  | -0.257 |  | 0.125 |  | -5.652 |  | -1.256 |  | -2.164 |  | -0.348 |  | < .001 | *** |
|  |  | SyncFirst, sync, hard, group incorrect |  | -1.207 |  | -1.752 |  | -0.662 |  | 0.152 |  | -7.941 |  | -2.145 |  | -3.368 |  | -0.921 |  | < .001 | *** |
|  |  | AsyncFirst, async, easy, group correct |  | -0.139 |  | -0.605 |  | 0.328 |  | 0.130 |  | -1.065 |  | -0.246 |  | -1.079 |  | 0.586 |  | 1.000 |  |
|  |  | SyncFirst, async, easy, group correct |  | -0.003 |  | -0.548 |  | 0.542 |  | 0.152 |  | -0.019 |  | -0.005 |  | -0.974 |  | 0.964 |  | 1.000 |  |
|  |  | AsyncFirst, sync, easy, group correct |  | 0.030 |  | -0.366 |  | 0.426 |  | 0.110 |  | 0.272 |  | 0.053 |  | -0.648 |  | 0.755 |  | 1.000 |  |
|  |  | SyncFirst, sync, easy, group correct |  | -0.163 |  | -0.708 |  | 0.382 |  | 0.152 |  | -1.071 |  | -0.289 |  | -1.263 |  | 0.685 |  | 1.000 |  |
|  |  | AsyncFirst, async, hard, group correct |  | -0.738 |  | -1.226 |  | -0.250 |  | 0.136 |  | -5.438 |  | -1.312 |  | -2.289 |  | -0.334 |  | < .001 | *** |
|  |  | SyncFirst, async, hard, group correct |  | -0.497 |  | -1.042 |  | 0.048 |  | 0.152 |  | -3.269 |  | -0.883 |  | -1.899 |  | 0.134 |  | 0.073 |  |
|  |  | AsyncFirst, sync, hard, group correct |  | -0.547 |  | -1.013 |  | -0.081 |  | 0.129 |  | -4.226 |  | -0.972 |  | -1.863 |  | -0.081 |  | 0.003 | ** |
|  |  | SyncFirst, sync, hard, group correct |  | -0.555 |  | -1.100 |  | -0.010 |  | 0.152 |  | -3.650 |  | -0.986 |  | -2.014 |  | 0.042 |  | 0.022 | * |
| SyncFirst, sync, easy, group incorrect |  | AsyncFirst, async, hard, group incorrect |  | -1.017 |  | -1.562 |  | -0.472 |  | 0.152 |  | -6.689 |  | -1.807 |  | -2.962 |  | -0.651 |  | < .001 | *** |
|  |  | SyncFirst, async, hard, group incorrect |  | -0.596 |  | -1.049 |  | -0.142 |  | 0.126 |  | -4.718 |  | -1.058 |  | -1.943 |  | -0.173 |  | < .001 | *** |
|  |  | AsyncFirst, sync, hard, group incorrect |  | -0.560 |  | -1.105 |  | -0.015 |  | 0.152 |  | -3.684 |  | -0.995 |  | -2.024 |  | 0.034 |  | 0.020 | * |
|  |  | SyncFirst, sync, hard, group incorrect |  | -1.060 |  | -1.486 |  | -0.634 |  | 0.118 |  | -8.954 |  | -1.884 |  | -2.884 |  | -0.884 |  | < .001 | *** |
|  |  | AsyncFirst, async, easy, group correct |  | 0.008 |  | -0.537 |  | 0.553 |  | 0.152 |  | 0.055 |  | 0.015 |  | -0.954 |  | 0.984 |  | 1.000 |  |
|  |  | SyncFirst, async, easy, group correct |  | 0.144 |  | -0.298 |  | 0.586 |  | 0.123 |  | 1.170 |  | 0.256 |  | -0.534 |  | 1.046 |  | 1.000 |  |
|  |  | AsyncFirst, sync, easy, group correct |  | 0.177 |  | -0.368 |  | 0.722 |  | 0.152 |  | 1.164 |  | 0.314 |  | -0.661 |  | 1.289 |  | 1.000 |  |
|  |  | SyncFirst, sync, easy, group correct |  | -0.016 |  | -0.391 |  | 0.359 |  | 0.104 |  | -0.152 |  | -0.028 |  | -0.693 |  | 0.637 |  | 1.000 |  |
|  |  | AsyncFirst, async, hard, group correct |  | -0.591 |  | -1.136 |  | -0.046 |  | 0.152 |  | -3.889 |  | -1.050 |  | -2.086 |  | -0.015 |  | 0.010 | ** |
|  |  | SyncFirst, async, hard, group correct |  | -0.350 |  | -0.812 |  | 0.112 |  | 0.128 |  | -2.723 |  | -0.622 |  | -1.469 |  | 0.226 |  | 0.370 |  |
|  |  | AsyncFirst, sync, hard, group correct |  | -0.400 |  | -0.945 |  | 0.145 |  | 0.152 |  | -2.633 |  | -0.711 |  | -1.711 |  | 0.289 |  | 0.445 |  |
|  |  | SyncFirst, sync, hard, group correct |  | -0.408 |  | -0.849 |  | 0.033 |  | 0.123 |  | -3.327 |  | -0.725 |  | -1.546 |  | 0.096 |  | 0.065 |  |
| AsyncFirst, async, hard, group incorrect |  | SyncFirst, async, hard, group incorrect |  | 0.421 |  | -0.124 |  | 0.966 |  | 0.152 |  | 2.771 |  | 0.748 |  | -0.255 |  | 1.752 |  | 0.328 |  |
|  |  | AsyncFirst, sync, hard, group incorrect |  | 0.457 |  | 0.064 |  | 0.849 |  | 0.109 |  | 4.190 |  | 0.812 |  | 0.062 |  | 1.561 |  | 0.004 | ** |
|  |  | SyncFirst, sync, hard, group incorrect |  | -0.043 |  | -0.588 |  | 0.502 |  | 0.152 |  | -0.285 |  | -0.077 |  | -1.046 |  | 0.892 |  | 1.000 |  |
|  |  | AsyncFirst, async, easy, group correct |  | 1.025 |  | 0.559 |  | 1.491 |  | 0.129 |  | 7.919 |  | 1.822 |  | 0.782 |  | 2.861 |  | < .001 | *** |
|  |  | SyncFirst, async, easy, group correct |  | 1.161 |  | 0.616 |  | 1.706 |  | 0.152 |  | 7.637 |  | 2.063 |  | 0.856 |  | 3.269 |  | < .001 | *** |
|  |  | AsyncFirst, sync, easy, group correct |  | 1.194 |  | 0.706 |  | 1.682 |  | 0.136 |  | 8.795 |  | 2.121 |  | 0.985 |  | 3.257 |  | < .001 | *** |
|  |  | SyncFirst, sync, easy, group correct |  | 1.001 |  | 0.456 |  | 1.546 |  | 0.152 |  | 6.585 |  | 1.778 |  | 0.628 |  | 2.929 |  | < .001 | *** |
|  |  | AsyncFirst, async, hard, group correct |  | 0.426 |  | 0.029 |  | 0.822 |  | 0.110 |  | 3.863 |  | 0.756 |  | 0.007 |  | 1.505 |  | 0.011 | * |
|  |  | SyncFirst, async, hard, group correct |  | 0.667 |  | 0.122 |  | 1.212 |  | 0.152 |  | 4.387 |  | 1.185 |  | 0.132 |  | 2.238 |  | 0.002 | ** |
|  |  | AsyncFirst, sync, hard, group correct |  | 0.617 |  | 0.150 |  | 1.083 |  | 0.130 |  | 4.739 |  | 1.096 |  | 0.184 |  | 2.007 |  | < .001 | *** |
|  |  | SyncFirst, sync, hard, group correct |  | 0.609 |  | 0.064 |  | 1.154 |  | 0.152 |  | 4.006 |  | 1.082 |  | 0.042 |  | 2.122 |  | 0.006 | ** |
| SyncFirst, async, hard, group incorrect |  | AsyncFirst, sync, hard, group incorrect |  | 0.036 |  | -0.509 |  | 0.581 |  | 0.152 |  | 0.235 |  | 0.063 |  | -0.906 |  | 1.033 |  | 1.000 |  |
|  |  | SyncFirst, sync, hard, group incorrect |  | -0.465 |  | -0.836 |  | -0.093 |  | 0.103 |  | -4.499 |  | -0.825 |  | -1.544 |  | -0.107 |  | 0.001 | ** |
|  |  | AsyncFirst, async, easy, group correct |  | 0.604 |  | 0.059 |  | 1.149 |  | 0.152 |  | 3.973 |  | 1.073 |  | 0.035 |  | 2.112 |  | 0.007 | ** |
|  |  | SyncFirst, async, easy, group correct |  | 0.740 |  | 0.299 |  | 1.181 |  | 0.123 |  | 6.034 |  | 1.314 |  | 0.409 |  | 2.220 |  | < .001 | *** |
|  |  | AsyncFirst, sync, easy, group correct |  | 0.772 |  | 0.228 |  | 1.317 |  | 0.152 |  | 5.082 |  | 1.373 |  | 0.292 |  | 2.453 |  | < .001 | *** |
|  |  | SyncFirst, sync, easy, group correct |  | 0.580 |  | 0.118 |  | 1.042 |  | 0.128 |  | 4.511 |  | 1.030 |  | 0.136 |  | 1.924 |  | 0.001 | ** |
|  |  | AsyncFirst, async, hard, group correct |  | 0.004 |  | -0.540 |  | 0.549 |  | 0.152 |  | 0.029 |  | 0.008 |  | -0.961 |  | 0.977 |  | 1.000 |  |
|  |  | SyncFirst, async, hard, group correct |  | 0.246 |  | -0.129 |  | 0.621 |  | 0.104 |  | 2.355 |  | 0.437 |  | -0.246 |  | 1.119 |  | 0.889 |  |
|  |  | AsyncFirst, sync, hard, group correct |  | 0.195 |  | -0.349 |  | 0.740 |  | 0.152 |  | 1.286 |  | 0.347 |  | -0.629 |  | 1.324 |  | 1.000 |  |
|  |  | SyncFirst, sync, hard, group correct |  | 0.188 |  | -0.254 |  | 0.630 |  | 0.123 |  | 1.524 |  | 0.334 |  | -0.460 |  | 1.127 |  | 1.000 |  |
| AsyncFirst, sync, hard, group incorrect |  | SyncFirst, sync, hard, group incorrect |  | -0.500 |  | -1.045 |  | 0.045 |  | 0.152 |  | -3.291 |  | -0.889 |  | -1.906 |  | 0.128 |  | 0.069 |  |
|  |  | AsyncFirst, async, easy, group correct |  | 0.568 |  | 0.080 |  | 1.056 |  | 0.136 |  | 4.187 |  | 1.010 |  | 0.077 |  | 1.943 |  | 0.004 | ** |
|  |  | SyncFirst, async, easy, group correct |  | 0.704 |  | 0.159 |  | 1.249 |  | 0.152 |  | 4.632 |  | 1.251 |  | 0.189 |  | 2.313 |  | < .001 | *** |
|  |  | AsyncFirst, sync, easy, group correct |  | 0.737 |  | 0.271 |  | 1.202 |  | 0.129 |  | 5.692 |  | 1.309 |  | 0.367 |  | 2.251 |  | < .001 | *** |
|  |  | SyncFirst, sync, easy, group correct |  | 0.544 |  | -8.998×10^-4^ |  | 1.089 |  | 0.152 |  | 3.579 |  | 0.967 |  | -0.059 |  | 1.992 |  | 0.028 | * |
|  |  | AsyncFirst, async, hard, group correct |  | -0.031 |  | -0.498 |  | 0.435 |  | 0.130 |  | -0.240 |  | -0.056 |  | -0.884 |  | 0.773 |  | 1.000 |  |
|  |  | SyncFirst, async, hard, group correct |  | 0.210 |  | -0.335 |  | 0.755 |  | 0.152 |  | 1.382 |  | 0.373 |  | -0.604 |  | 1.351 |  | 1.000 |  |
|  |  | AsyncFirst, sync, hard, group correct |  | 0.160 |  | -0.236 |  | 0.556 |  | 0.110 |  | 1.450 |  | 0.284 |  | -0.424 |  | 0.992 |  | 1.000 |  |
|  |  | SyncFirst, sync, hard, group correct |  | 0.152 |  | -0.393 |  | 0.697 |  | 0.152 |  | 1.000 |  | 0.270 |  | -0.703 |  | 1.244 |  | 1.000 |  |
| SyncFirst, sync, hard, group incorrect |  | AsyncFirst, async, easy, group correct |  | 1.068 |  | 0.524 |  | 1.613 |  | 0.152 |  | 7.030 |  | 1.899 |  | 0.725 |  | 3.072 |  | < .001 | *** |
|  |  | SyncFirst, async, easy, group correct |  | 1.204 |  | 0.742 |  | 1.666 |  | 0.128 |  | 9.371 |  | 2.140 |  | 1.032 |  | 3.247 |  | < .001 | *** |
|  |  | AsyncFirst, sync, easy, group correct |  | 1.237 |  | 0.692 |  | 1.782 |  | 0.152 |  | 8.139 |  | 2.198 |  | 0.963 |  | 3.433 |  | < .001 | *** |
|  |  | SyncFirst, sync, easy, group correct |  | 1.044 |  | 0.603 |  | 1.485 |  | 0.123 |  | 8.519 |  | 1.856 |  | 0.841 |  | 2.870 |  | < .001 | *** |
|  |  | AsyncFirst, async, hard, group correct |  | 0.469 |  | -0.076 |  | 1.014 |  | 0.152 |  | 3.085 |  | 0.833 |  | -0.178 |  | 1.845 |  | 0.130 |  |
|  |  | SyncFirst, async, hard, group correct |  | 0.710 |  | 0.268 |  | 1.152 |  | 0.123 |  | 5.766 |  | 1.262 |  | 0.362 |  | 2.162 |  | < .001 | *** |
|  |  | AsyncFirst, sync, hard, group correct |  | 0.660 |  | 0.115 |  | 1.205 |  | 0.152 |  | 4.342 |  | 1.173 |  | 0.121 |  | 2.224 |  | 0.002 | ** |
|  |  | SyncFirst, sync, hard, group correct |  | 0.652 |  | 0.277 |  | 1.027 |  | 0.104 |  | 6.254 |  | 1.159 |  | 0.381 |  | 1.937 |  | < .001 | *** |
| AsyncFirst, async, easy, group correct |  | SyncFirst, async, easy, group correct |  | 0.136 |  | -0.409 |  | 0.681 |  | 0.152 |  | 0.893 |  | 0.241 |  | -0.731 |  | 1.214 |  | 1.000 |  |
|  |  | AsyncFirst, sync, easy, group correct |  | 0.169 |  | -0.224 |  | 0.561 |  | 0.109 |  | 1.546 |  | 0.300 |  | -0.403 |  | 1.002 |  | 1.000 |  |
|  |  | SyncFirst, sync, easy, group correct |  | -0.024 |  | -0.569 |  | 0.521 |  | 0.152 |  | -0.159 |  | -0.043 |  | -1.012 |  | 0.926 |  | 1.000 |  |
|  |  | AsyncFirst, async, hard, group correct |  | -0.599 |  | -1.050 |  | -0.149 |  | 0.125 |  | -4.794 |  | -1.065 |  | -1.944 |  | -0.187 |  | < .001 | *** |
|  |  | SyncFirst, async, hard, group correct |  | -0.358 |  | -0.903 |  | 0.187 |  | 0.152 |  | -2.357 |  | -0.637 |  | -1.630 |  | 0.357 |  | 0.889 |  |
|  |  | AsyncFirst, sync, hard, group correct |  | -0.408 |  | -0.888 |  | 0.071 |  | 0.133 |  | -3.064 |  | -0.726 |  | -1.612 |  | 0.160 |  | 0.138 |  |
|  |  | SyncFirst, sync, hard, group correct |  | -0.416 |  | -0.961 |  | 0.129 |  | 0.152 |  | -2.738 |  | -0.740 |  | -1.742 |  | 0.263 |  | 0.355 |  |
| SyncFirst, async, easy, group correct |  | AsyncFirst, sync, easy, group correct |  | 0.033 |  | -0.512 |  | 0.578 |  | 0.152 |  | 0.216 |  | 0.058 |  | -0.911 |  | 1.027 |  | 1.000 |  |
|  |  | SyncFirst, sync, easy, group correct |  | -0.160 |  | -0.531 |  | 0.212 |  | 0.103 |  | -1.549 |  | -0.284 |  | -0.950 |  | 0.381 |  | 1.000 |  |
|  |  | AsyncFirst, async, hard, group correct |  | -0.735 |  | -1.280 |  | -0.190 |  | 0.152 |  | -4.837 |  | -1.306 |  | -2.377 |  | -0.236 |  | < .001 | *** |
|  |  | SyncFirst, async, hard, group correct |  | -0.494 |  | -0.920 |  | -0.068 |  | 0.118 |  | -4.172 |  | -0.878 |  | -1.692 |  | -0.063 |  | 0.004 | ** |
|  |  | AsyncFirst, sync, hard, group correct |  | -0.544 |  | -1.089 |  | 6.805×10^-4^ |  | 0.152 |  | -3.581 |  | -0.967 |  | -1.993 |  | 0.059 |  | 0.028 | * |
|  |  | SyncFirst, sync, hard, group correct |  | -0.552 |  | -1.006 |  | -0.098 |  | 0.126 |  | -4.372 |  | -0.981 |  | -1.855 |  | -0.106 |  | 0.002 | ** |
| AsyncFirst, sync, easy, group correct |  | SyncFirst, sync, easy, group correct |  | -0.193 |  | -0.738 |  | 0.352 |  | 0.152 |  | -1.268 |  | -0.343 |  | -1.319 |  | 0.634 |  | 1.000 |  |
|  |  | AsyncFirst, async, hard, group correct |  | -0.768 |  | -1.247 |  | -0.289 |  | 0.133 |  | -5.761 |  | -1.365 |  | -2.337 |  | -0.392 |  | < .001 | *** |
|  |  | SyncFirst, async, hard, group correct |  | -0.527 |  | -1.072 |  | 0.018 |  | 0.152 |  | -3.466 |  | -0.936 |  | -1.958 |  | 0.086 |  | 0.040 | * |
|  |  | AsyncFirst, sync, hard, group correct |  | -0.577 |  | -1.027 |  | -0.127 |  | 0.125 |  | -4.615 |  | -1.025 |  | -1.898 |  | -0.153 |  | < .001 | *** |
|  |  | SyncFirst, sync, hard, group correct |  | -0.585 |  | -1.130 |  | -0.040 |  | 0.152 |  | -3.847 |  | -1.039 |  | -2.073 |  | -0.005 |  | 0.011 | * |
| SyncFirst, sync, easy, group correct |  | AsyncFirst, async, hard, group correct |  | -0.575 |  | -1.120 |  | -0.030 |  | 0.152 |  | -3.785 |  | -1.022 |  | -2.055 |  | 0.010 |  | 0.014 | * |
|  |  | SyncFirst, async, hard, group correct |  | -0.334 |  | -0.788 |  | 0.120 |  | 0.126 |  | -2.646 |  | -0.594 |  | -1.424 |  | 0.237 |  | 0.443 |  |
|  |  | AsyncFirst, sync, hard, group correct |  | -0.384 |  | -0.929 |  | 0.161 |  | 0.152 |  | -2.528 |  | -0.683 |  | -1.680 |  | 0.315 |  | 0.584 |  |
|  |  | SyncFirst, sync, hard, group correct |  | -0.392 |  | -0.818 |  | 0.034 |  | 0.118 |  | -3.310 |  | -0.696 |  | -1.489 |  | 0.096 |  | 0.068 |  |
| AsyncFirst, async, hard, group correct |  | SyncFirst, async, hard, group correct |  | 0.241 |  | -0.304 |  | 0.786 |  | 0.152 |  | 1.587 |  | 0.429 |  | -0.552 |  | 1.409 |  | 1.000 |  |
|  |  | AsyncFirst, sync, hard, group correct |  | 0.191 |  | -0.201 |  | 0.583 |  | 0.109 |  | 1.752 |  | 0.339 |  | -0.365 |  | 1.044 |  | 1.000 |  |
|  |  | SyncFirst, sync, hard, group correct |  | 0.183 |  | -0.362 |  | 0.728 |  | 0.152 |  | 1.206 |  | 0.326 |  | -0.650 |  | 1.301 |  | 1.000 |  |
| SyncFirst, async, hard, group correct |  | AsyncFirst, sync, hard, group correct |  | -0.050 |  | -0.595 |  | 0.495 |  | 0.152 |  | -0.331 |  | -0.089 |  | -1.059 |  | 0.880 |  | 1.000 |  |
|  |  | SyncFirst, sync, hard, group correct |  | -0.058 |  | -0.429 |  | 0.314 |  | 0.103 |  | -0.561 |  | -0.103 |  | -0.762 |  | 0.556 |  | 1.000 |  |
| AsyncFirst, sync, hard, group correct |  | SyncFirst, sync, hard, group correct |  | -0.008 |  | -0.553 |  | 0.537 |  | 0.152 |  | -0.050 |  | -0.014 |  | -0.983 |  | 0.955 |  | 1.000 |  |
| * p < .05, ** p < .01, *** p < .001 | | | | | | | | | | | | | | | | | | | | | |
| *Note.*  P-value and confidence intervals adjusted for comparing a family of 120 estimates (confidence intervals corrected using the bonferroni method). | | | | | | | | | | | | | | | | | | | | | |
